# Quasistable multiclonal carcinomas evolve through mutations with locally extended phenotypes

**DOI:** 10.64898/2026.11.13.766058

**Authors:** Mst Shanta Khatun, Erwin P. Lobo, Mary R. Myerscough, Peter S. Kim, J. Guy Lyons

## Abstract

The earliest stages of epithelial carcinogenesis are often considered to arise via lineal evolution, a mutation-driven process characterised by purely competitive clonal interactions. However, this conventional view cannot fully explain the persistent polyclonality observed in many cancers. We therefore investigated an alternative mechanism of cancer progression, termed interclonal cooperation, arising from mutations that cause cells to produce microenvironmental changes, enabling them to affect the behaviours of surrounding cells. We built a Voronoi cell-based model (VCBM) of epithelial carcinogenesis evolution to examine the role of cancer cell interactions in proliferation, motility and degradation of the extracellular matrix (ECM) during the earliest stages of carcinogenesis. In this model, we considered mutations with four distinct effects on cells: impaired apoptosis, faster cell-cycling, hyperproliferation and enhanced invasiveness. In each case, these phenotypes can be either cell-autonomous, affecting only the mutated cell itself, or microenvironmental, affecting both the mutated cell and the surrounding cells within a specified range. Using our model, we investigated how microenvironmentally extended phenotypes in epithelial cells influence the formation of carcinogenic lesions. We found that the acquisition of microenvironmentally extended phenotypes reduces the overall mutational burden of the tissue by increasing the competitiveness of non-mutant cells with cells harbouring cell-autonomous mutations. However, progression from pre-malignancy to malignancy can occur without the acquisition of additional mutations, through field collision, i.e. by the growth and/or migration of mutually beneficial clones, leading to their co-localisation, which enables malignant interactions. Our results also indicate that mutations with microenvironmentally extended phenotypes can enhance the speed with which malignancy occurs compared with cell-autonomous phenotypes, through symbiotic clonal interactions. An important contribution to the speed of carcinogenesis is the ability of microenvironmentally extended phenotypes to overcome mutual inhibition of pro-malignant intracellular signalling pathways. Moreover, we found that, after an initial period of rapid change and clonal succession during carcinogenesis, genetically distinct clones of cell types persist together for long periods, maintaining stable proportions.

**Author summary:** Cancer progression is regulated not only by genetic mutations within individual cells, but also by interactions between genetically different cancer cell populations. Many models of cancer focus on competitive single-cell behaviour, whereas few studies have investigated how cooperative interactions among cancer cell clones can affect the early stages of carcinogenesis. To this end, we developed a computational model to investigate how cooperative cell-cell interactions influence the malignant behaviour of cells during the early stages of epithelial carcinogenesis. Our results show that cooperative cell populations can become malignant faster than non-cooperative cells, especially in cases where oncogenic signalling pathways are mutually incompatible within individual cells, and result in persistent clonal heterogeneity within the malignant lesion. These findings highlight the importance of cell-cell interactions in the early stages of tumour growth that progress to cancer and demonstrate how mathematical modelling can provide insight into the complex evolutionary processes of cancer, which are challenging to observe experimentally.

## 1 Introduction

Carcinogenesis is an evolutionary process in which cells acquire random mutations, giving rise to clones of cells with a common genetic composition. These mutations cause unregulated tissue growth through aggressive cell proliferation, impaired cell death, increased motility and invasion, and metastasis. A common paradigm postulates that tumours evolve through clonal competition: clones of cells whose mutations confer an advantage to their replication will outcompete clones that lack the mutation or have neutral or deleterious mutations. In this process, which we call lineal evolution, cancer arises from a single cell that has acquired all of the pro-malignant mutations required to maintain malignancy [1, 2]. However, many cancers are genetically heterogeneous, and a high level of intratumoral heterogeneity (ITH) predicts poor patient outcomes [3, 4]. During early carcinogenesis, a transient ITH may be caused by a lag in the domination of the tissue by the fittest clone during lineal evolution. However, interactions between cancer cells can also contribute to ITH through interclonal co-operativity, in which polyclonal cancers evolve symbiotically, either from a single cell whose mutated clonal progeny interact and cooperate with other derivative clones, or from clones derived from two or more nearby cells whose mutated offspring interact and cooperate with each other for the progression of tumour [5–14]. Interclonal co-operativity can occur when mutations cause cells to produce phenotypes that extend to their microenvironments, influencing the behaviours of cells around them, but little investigation has been done to investigate the qualitative and quantitative impact on this mode of progression compared with lineal evolution.

Carcinomas are the most common type of cancer and arise from epithelial tissues, in which the cells are spatially constrained laterally, by direct contact with neighbouring cells, and vertically, by the underlying ECM. To be considered malignant, the cells must both proliferate abnormally quickly and invade the ECM. Spatial constraints can impact the pattern of evolution in unpredictable ways [15]. In the case of cancer cells and their precursors, the spatial constraints can lead to field cancerisation, in which mutated cells form spatially contiguous pre-malignant “fields” that appear to be clinically normal but nevertheless have a survival or proliferative advantage over non-mutant cells and have a high probability of progressing to malignancy [16–18]. Key mutations on the path to the evolution of epithelial carcinogenesis reduce a cell’s death rate, increase proliferation, and increase invasion [19]. In the case of head and neck squamous cell carcinomas (HNSCCs), examples of genetic alterations that epithelial cells acquire successively during oral carcinogenesis include *TP53* mutations, which cause an increase in cell lifespan by blocking programmed cell death [20], *NOTCH* mutations, which cause cells to acquire a greater replication capacity by disrupting normal control of cell differentiation [21], extension of telomeres, leading to a higher replication potential to the cells [22], *HRAS* mutations, which activate mitogen-activated kinase pathways leading to hyperproliferation [7, 23–26], and increased *SNAI2* expression, which enhances motility and ECM invasion [23, 27, 28] (Figure 1).

**Figure 1:**
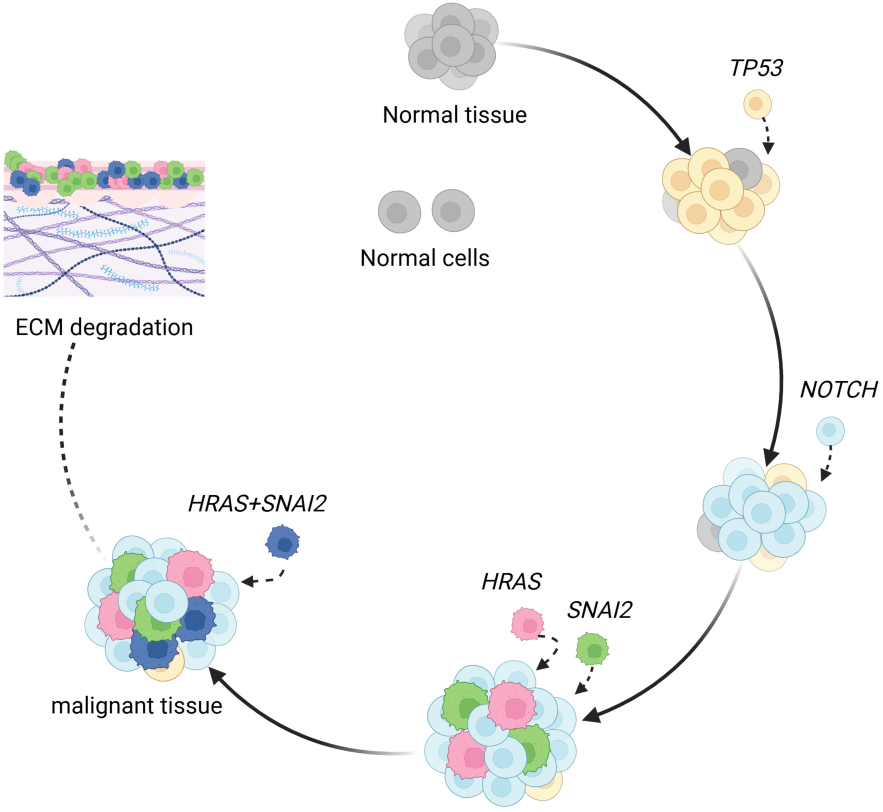
HNSCC evolution as an example of epithelial cancer evolution. Normal cells first acquire *TP53* mutations, then these *TP53* mutated cells acquire *NOTCH* mutations. Then, these cells acquire mutations that either activate *HRAS* or increase expression of *SNAI2* in either order. *HRAS* mutant cells are hyperproliferative and induce hyperproliferation in nearby cells. *SNAI2* expressing cells undergo an epithelial-mesenchymal transition, resulting in higher motility and the induction of higher motility in nearby cells, and also respond to nearby HRAS cells by degrading ECM. Figure created with BioRender.com (https://BioRender.com/4ps62hc).

Mathematical modelling has been widely used to investigate the evolution of cancer [29–36]. Agent-based models (ABMs), in particular, enable the explicit representation of the proliferation, death and movement of individual cells and their local interactions, providing a powerful framework for investigating tumour evolution. Furthermore, several ABMs incorporate ECM structure and cell-ECM interactions, enabling investigation of how ECM mechanics and remodelling influence cell migration and invasion within tissue environments [37, 38].

Within the class of ABMs, lattice-free frameworks allow individual cells to move and interact in continuous space, enabling a more realistic representation of cell morphology, mechanical interactions, and spatial rearrangements than cellular automata [39–42]. For instance, active finite lattice-free VCBMs have been used to study collective cell motility and epithelial-mesenchymal transitions in confluent epithelial tissues [43]. Similarly, multi-scale lattice-free ABMs have been applied to investigate early tumour growth while accounting for cell-cell interactions within epithelial tissue [44]. By capturing emergent collective behaviour and incorporating cell-cell and cell-matrix interactions, ABMs offer mechanistic insight into the processes driving cancer progression [45–49].

In the present study, we extended the work of Lobo [8] and built a VCBM to investigate the early stages of epithelial cell evolution that lead to cancer. In this VCBM, we incorporated the ECM, cellular genetic heterogeneity, cell-cell interactions, cell-ECM interactions and probabilistic variability. Building on this model, we focused on mutations that have microenvironmental phenotypes and their effects in forming a heterogeneous, cooperative malignant lesion that progresses to cancer. We investigated and compared mutations having cell-autonomous phenotypes with those having microenvironmental phenotypes, their relationships to lineal evolution and interclonal cooperativity, and explored which of these evolutionary mechanisms is more likely to produce a malignant lesion. Our results indicate that mutations with microenvironmentally extended phenotypes can accelerate malignant progression compared to cell-autonomous phenotypes through symbiotic clonal interactions. Furthermore, progression from pre-malignancy to malignancy can occur without the acquisition of additional mutations, through field collision, i.e. by the growth and/or migration of mutually beneficial clones, leading to their co-localisation, which enables malignant interactions. In addition, after a period of rapid change and clonal succession during carcinogenesis, genetically distinct clones of cell types persist together for long periods, maintaining stable proportions. Finally, microenvironmental phenotypes can overcome the limits on carcinogenesis caused by mutual inhibition of pro-malignant intracellular signalling pathways.

## 2 Model description

### 2.1 Voronoi cell-based model

Agent-based models have become increasingly popular as they provide a convenient framework for incorporating cell heterogeneity, spatial information and probabilistic interactions. They come in a variety of forms, including lattice-based, lattice-free, and hybrid discrete-continuum models. In this study, we develop a lattice-free 2-D VCBM for modelling interactions between genetically distinct clones during the earliest stages of cancer development [8, 39, 41, 50–52], where we assume that spring forces exist between cells and cell locations are updated by solving an equation of motion [40, 51]. The Voronoi diagram resembles an epithelial cell monolayer, such as those of simple epithelial tissues or the basal layer of stratified epithelial tissues, which give rise to carcinomas, the most common forms of cancer. Our specific interest is to investigate how the microenvironmental effects of mutations in epithelial cell clones propel their progression to malignancy.

In our VCBM, the position of each cell is specified by points in 2-D space known as Voronoi generators or seeds. Each Voronoi generator corresponds to a Voronoi cell or polygon, which comprises the collection of points that are closer to that Voronoi generator than any other (Figure 2a). Cells sharing common edges are neighbours (Figure 2b), and cells move due to forces from neighbouring cells. Cells also proliferate and die with certain probabilities. The collection of Voronoi cells determines the Voronoi diagram or tessellation (Figure 2c). For our boundary, we set an outer rim of cells that do not move, proliferate, or die and form a buffer on which other cells have no effect, giving us a contained region of tissue.

**Figure 2:**
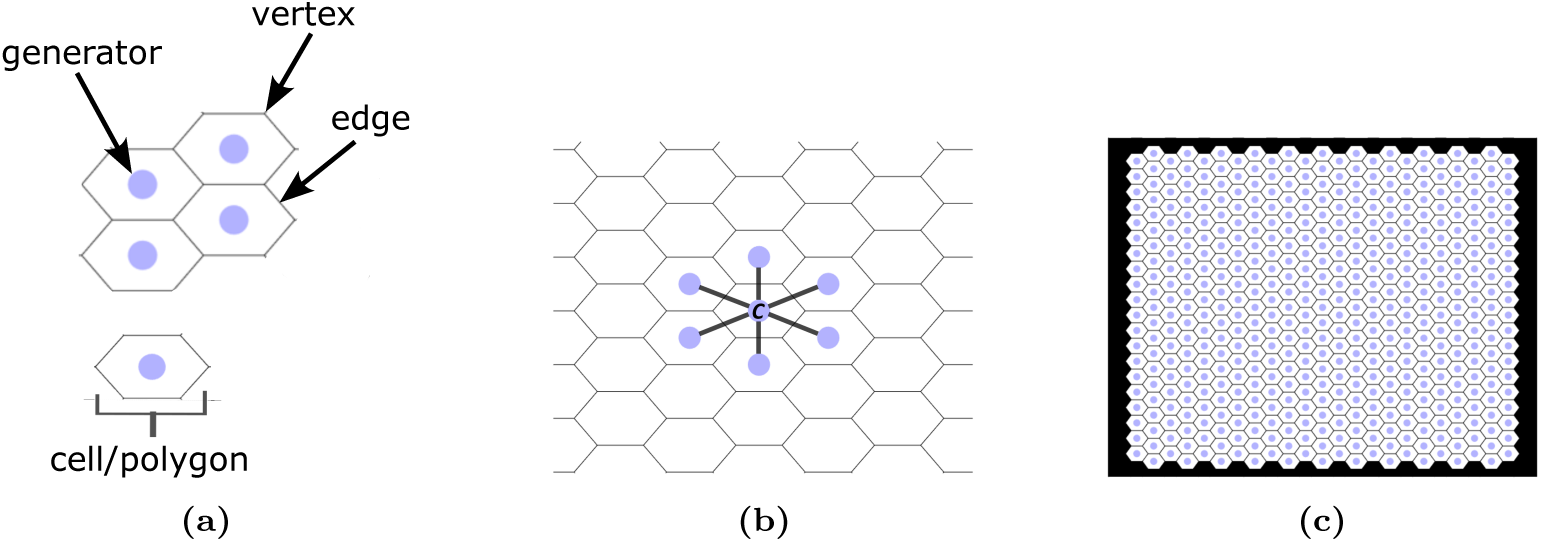
(a) Voronoi generators, edges, vertices, a cell/polygon. (b) Neighbours of Voronoi cell *C*. (c) Representation of Voronoi tessellation with outer boundary. In the illustration, light purple circles represent the Voronoi centres, and these Voronoi centres or seeds with boundaries represent Voronoi cells. The black border on the four sides represents the boundary, formed by a rim of cells that do not move, proliferate, or die.

A flow chart of the VCBM is presented in Figure 3. Following [8, 40, 53], we designed the initial seeds in the lattice in such a way that the corresponding Voronoi cells make a hexagonal tessellation. Delaunay triangulations are used to determine the neighbours of a particular cell in the Voronoi tessellation, and details of this procedure are illustrated in [39–41, 52–54].

**Figure 3:**
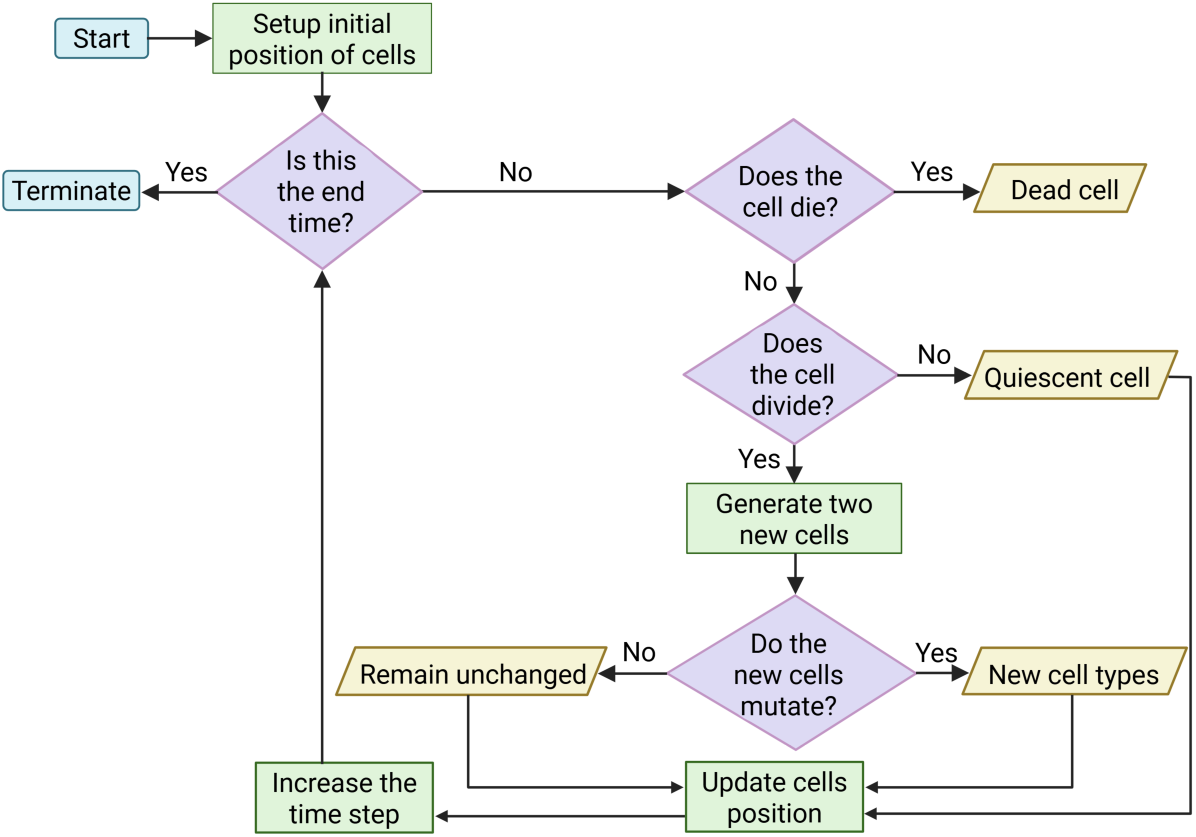
Flow chart of the Voronoi Cell-based computational model algorithm. Flow chart created with BioRender.com (https://BioRender.com/8eclpl0).

At each time step Δ*t*, cells undergo fundamental procedures such as movement, proliferation and death. In the following sections, we will discuss how we model those processes in the VCBM.

#### Cell movement

We consider cell movement due to the pressure created by neighbouring cells. We model this pressure by assuming that adjacent cells are connected by springs between their Voronoi centres (Figure 4). For simplicity, we assume the same spring rest length between each pair of cells and use Hooke’s law to calculate the net force on each cell from all neighbouring cells and then determine the displacement of each cell in the next time step.

**Figure 4:**
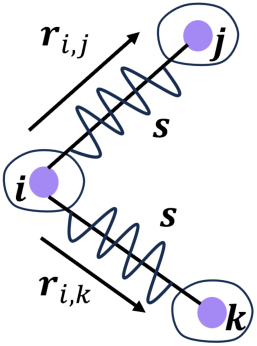
Springs connect cells *j* and *k* to cell *i*. In the illustration, the spring rest lengths, *s*, of the springs are the same.

We assume that two types of forces govern the movement of each cell: (a) interaction forces, modelled as a network of damped springs connecting a particular cell *i* to its neighbouring cells (which can be determined by Delaunay triangulation), and (b) viscous forces. Now, the displacement of cell *i* is given by the following equation of motion:

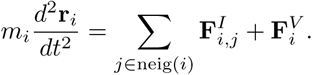

In the above equation, *m_i_* is the mass and ***r****_i_* is the position vector of cell *i*’s Voronoi centre, 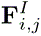 is the interaction force on cell *i* from neighbouring cell *j*, and 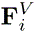 is the viscous force acting on cell *i*. The total interaction force 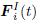, acting on the *i*th cell at time *t*, is equal to the sum of all the interaction forces coming from all the neighbouring cells *j ∈* neig(*i*) connected to cell *i* at that time and is given by the sum of spring forces from Hooke’s law:

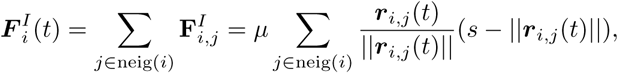

where *µ* is the spring constant, *||**r**_i,j_*(*t*)*||* is the Euclidean norm of vector ***r****_i,j_*(*t*) between cells *i* and *j* (see Figure 4), and *s* is the spring rest length [39–41, 53, 55]. Furthermore, we assume the cells are in a relatively dissipative environment, so motion can be approximated as being over-damped owing to strong friction, i.e. 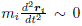 [39–
41, 53]. Also, the viscous force 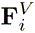 acting on cell *i* is proportional to its velocity, so

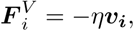

where ***v_i_*** is the velocity of cell *i* and *η* is the proportionality constant, known as the damping constant. The total force acting on cell *i* is given by

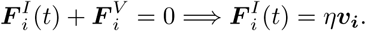

For a small time interval Δ*t*, the effective displacement of cell *i* is

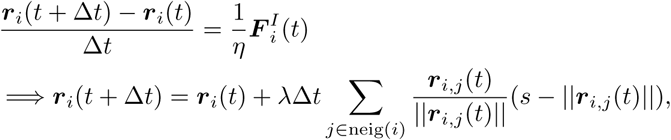

where *λ_d_* = *µ/η* is a cell motility parameter. In the model, we assume that the four different cell types *V* + *W*, *X*, *Y* and *X* + *Y* (see Section 2.1) move at varying motility rates *λ_V_* _+_*_W_*, *λ_X_*, *λ_Y_*, and *λ_X_*_+_*_Y_*, respectively.

#### Cell proliferation

At each time step Δ*t*, a cell will divide with probability

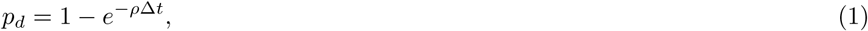

where *ρ* is the cell division rate, if

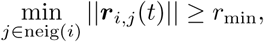

where *r*_min_ is the minimum distance required between a cell *i* and its neighbours for it to divide. When a cell *i* divides, a new Voronoi centre *k* is placed at a distance *d* away from the centre of the dividing cell in a random direction. Thus, *i* and *k* represent the two daughter cells in the Voronoi diagram.

We assume different cell types *V* + *W*, *X*, *Y*, and *X* + *Y* (see Section 2.1) proliferate at different rates *ρ_V_* _+_*_W_*, *ρ_X_*, *ρ_Y_*, and *ρ_X_*_+_*_Y_*, leading to corresponding probabilities *p_d,V_* _+_*_W_*, *p_d,X_*, *p_d,Y_*, and *p_d,X_*_+_*_Y_*, given by (1). During division, both the two daughter cells are equally likely to undergo mutation. If the original cell is of type *V* + *W*, it mutates to type *X* or *Y* with probability *ε*. If the original cell is an *X* or *Y* cell, it acquires a further mutation with the same probability *ε*, turning into an *X* + *Y* cell. In our model, we assume either or both daughter cells can acquire mutation during division.

#### Cell death

At each time step Δ*t*, each cell can die with probability

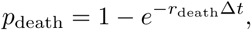

where *r*_death_ is the death rate. For simplicity, we assume all cells have the same death rate. Furthermore, when a cell dies, we remove it from the system as if it is consumed by neighbouring cells or delaminates into an upper layer [8].

#### Mutations and the phenotypes that they confer on cells

To model the evolution of epithelial carcinogenesis, we consider four types of mutations that epithelial cells successively acquire to reach malignancy. We call them *V_G_*, *W_G_*, *X_G_* and *Y_G_*. Of these, the first two mutations, *V_G_* and *W_G_*, always occur first and give rise to cells that have a longer replicative lifespan and a higher mutation rate than normal cells, but are not hyperproliferative or invasive. They also confer resistance to potential programmed cell death, such as might be induced by a subsequent oncogenic mutation. These phenotypes could be conferred, for example, by inactivating mutations in keratinocytes of *NOTCH* and *TP53* [56], respectively. The cells arising with *V_G_* and *W_G_* mutations would be predicted to outcompete normal cells in the tissue ecosystem. Nonetheless, these cells are still clinically undetectable and not malignant, but would create a pre-malignant field that serves as the basis for our simulations [8]. In addition, mutation *X_G_* confers more aggressive proliferation, and mutation *Y_G_* confers more invasiveness through both higher motility and enhanced ability to degrade ECM. For example, in keratinocytes, *HRAS* mutations confer hyperproliferation and mutations that increase expression of *SNAI2* induce an epithelial-mesenchymal transition (EMT) and thereby increase motility and ECM degradation [28].

#### Cell types

In our model, we consider four types of cells: (i) *V* +*W* cells (or pre-malignant cells), (ii) *X* cells (or hyperproliferative cells), (iii) *Y* cells (or invasive cells), and (iv) *X* + *Y* cells (both hyperproliferative and invasive cells). Herein, normal cells are not considered; all of the initial cells have acquired the first and second mutations (*V_G_*and *W_G_*) to give rise to *V* +*W* cells. To model a carcinoma that originates in a pre-malignant field, as typically happens in epithelial tissues [57], we generally use all *V* + *W* cells as an initial arrangement of cells in the Voronoi diagram. As these cells function similarly to healthy epithelial cells with respect to proliferation and invasiveness, except that they have a longer replicative lifespan, they are likely to dominate a patch of tissue before further mutations arise, forming a pre-malignant field. Then, these *V* + *W* cells can acquire either *X_G_* or *Y_G_* mutations with probability *ε*. Although the resulting cells contain 3 mutations, for simplicity, we refer to them as just *X* cells or *Y* cells, respectively. *X* cells and *Y* cells can subsequently acquire a further mutation with the same probability, producing *X* + *Y* cells expressing both mutations.

We have previously shown that cancer cells with a stable EMT phenotype depend on diffusible factors made by nearby hyperproliferative epithelial cells in order to proliferate rapidly [6]. As well as enhanced growth, the EMT cells respond to the epithelial cells by producing higher levels of proteinases that can diffuse over a short range and degrade the underlying ECM. Therefore, in our model, we assume invasive *Y* cells within a certain distance of nearby *X* or *X* + *Y* cells secrete ECM-degrading enzymes that diffuse over a short range and degrade the underlying ECM. We assume that *X* +*Y* cells secrete ECM-degrading enzymes as well, since they possess both mutations.

#### ECM degradation

To model ECM, we use a 36 *×* 36 square lattice that underlies the Voronoi tessellation representing cells. This square lattice has the same physical size as the Voronoi tessellation, and we let Δ*x* be the distance between adjacent points on the lattice. We assume that ECM is normally produced at a constant rate and has a maximal carrying capacity that we set to 1.

In addition, we assume that at each time step, the *Y* cells that are within a distance *r_x_*of either nearby *X* or *X* + *Y* cells, and the *X* + *Y* cells secrete ECM-degrading enzymes that diffuse over a short range and degrade the underlying ECM [6]. For simplicity, we assume that the secreting cell degrades the ECM within a fixed radius *r*_deg_ at a rate *σ_d_*.

Hence, the ECM production and degradation at point (*x, y*) in time Δ*t* is modelled by

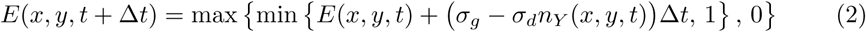

where *E*(*x, y, t*) is the ECM level at point (*x, y*) at time *t*, *σ_g_* is the ECM production rate, *σ_d_*is the ECM-degradation rate per enzyme-secreting cell, and *n_Y_* (*x, y, t*) is the number of enzyme secreting cells within radius *r*_deg_ of (*x, y*) at time *t*. Specifically, *n_Y_* (*x, y, t*) is the total number of enzyme-secreting *Y* (which are *Y* cells within *r_x_* of either *X* or *X* + *Y* cells) and *X* + *Y* cells whose Voronoi centres fall within the radius *r*_deg_ of (*x, y*) at time *t*.

#### Autonomous and microenvironmental phenotypes

For *X_G_*and *Y_G_*, we consider that they can produce one of two kinds of phenotypes: cell-autonomous (denoted by *a*), which impacts only the cell bearing this mutation but has no direct effect on the cells around it, or microenvironmental (denoted by *m*), which has an impact on both the cell bearing this mutation and the cells surrounding it [7]. In our model, the first two mutations, *V_G_* and *W_G_*, always give rise to cell-autonomous phenotypes, meaning that they do not interact with nearby cells. So, we denote the cells acquiring these first two mutations as *V_a_*+ *W_a_*. In contrast, the *X_G_* and *Y_G_* mutations can produce either cell-autonomous or microenvironmental phenotypes [8], which we denote *X_a_*, *X_m_*, *Y_a_*, or *Y_m_*. Hence, in the model, *X_m_* cells have an *X_G_* mutation that causes a hyperproliferative phenotype, and they confer this property on both themselves and their nearby cells. An example of *X_m_*is a mutation that causes the secretion of a diffusible growth factor. Similarly, *Y_m_* cells confer their invasive property on both themselves and nearby cells. An example of *Y_m_* is a mutation that causes the secretion of a cytokine that enhances motility and the production of ECM-degrading enzymes. In contrast, *X_a_* and *Y_a_* cells confer hyperproliferative and invasive properties on themselves, but not on their nearby cells.

To model the microenvironmental phenotype, we assume that a cell with an *X_m_*mutation influences all other cells within a distance *r* that lack *X_G_* mutations by increasing their proliferation probability to

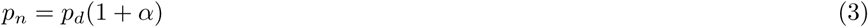

where *p_d_*is the basal proliferation probability rate and *α* is the strength by which the proliferation probability of nearby cells increases. Therefore, at each time step Δ*t*, all cells that are within *r* cell diameters of any *X_m_* cell will proliferate with an increased probability of *p_n_*, given in (3). For simplicity, we assume that the microenvironmental effects are at a saturating level and are neither additive nor synergistic. Thus, whether a cell is within *r* cell diameters of one *X_m_*cell or more than one *X_m_*cell, the strength by which proliferation increases is *α*.

Similarly, all cells that are within *r* cell diameters of one or more *Y_m_* cells will move at an increased motility rate *λ_n_*, given in (4):

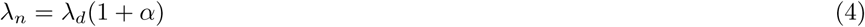

where *λ_d_*is the basal motility rate and *α* is the same strength by which the motility rate of nearby cells also increases.

In the case of ECM degradation, we assume that only the cells that have undergone an EMT (i.e. contain a *Y_a_* or *Y_m_* mutation) are able to degrade ECM [5–7, 23]. We assume the *Y* cells that have microenvironmental *X* or *X* + *Y* cells in their proximity (within a distance *r_x_*) degrade the underlying ECM at an increased rate to

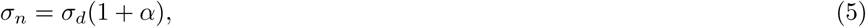

where *σ_d_*is the basal ECM-degradation rate and *α* is the strength at which the degradation rate increases—the same strength of microenvironmental influence at which the proliferation probability or motility rate of nearby cells increases.

#### Evolutionary trajectories

We considered four 2-step evolutionary scenarios in which each of the *X_G_* and *Y_G_* mutations was restricted to being either only cell autonomous or only microenvironmental (Figures 5A-D). Of these, in one scenario (Figure 5A), cells can become malignant only by two lineal evolution trajectories. In the other three scenarios, cells can become malignant either by lineal evolution trajectories or by symbiotic interactions mediated by microenvironmental phenotypes: commensal in Figure 5B and Figure 5C or cooperative in Figure 5D, and these three scenarios denote interclonal cooperativity. We identify these scenarios as Scenarios XaYa, XaYm, XmYa, and XmYm, depending on whether the *X_G_*and *Y_G_*mutations arise from cell-autonomous (*a*) or microenvironmental (*m*) phenotypes.

**Figure 5:**
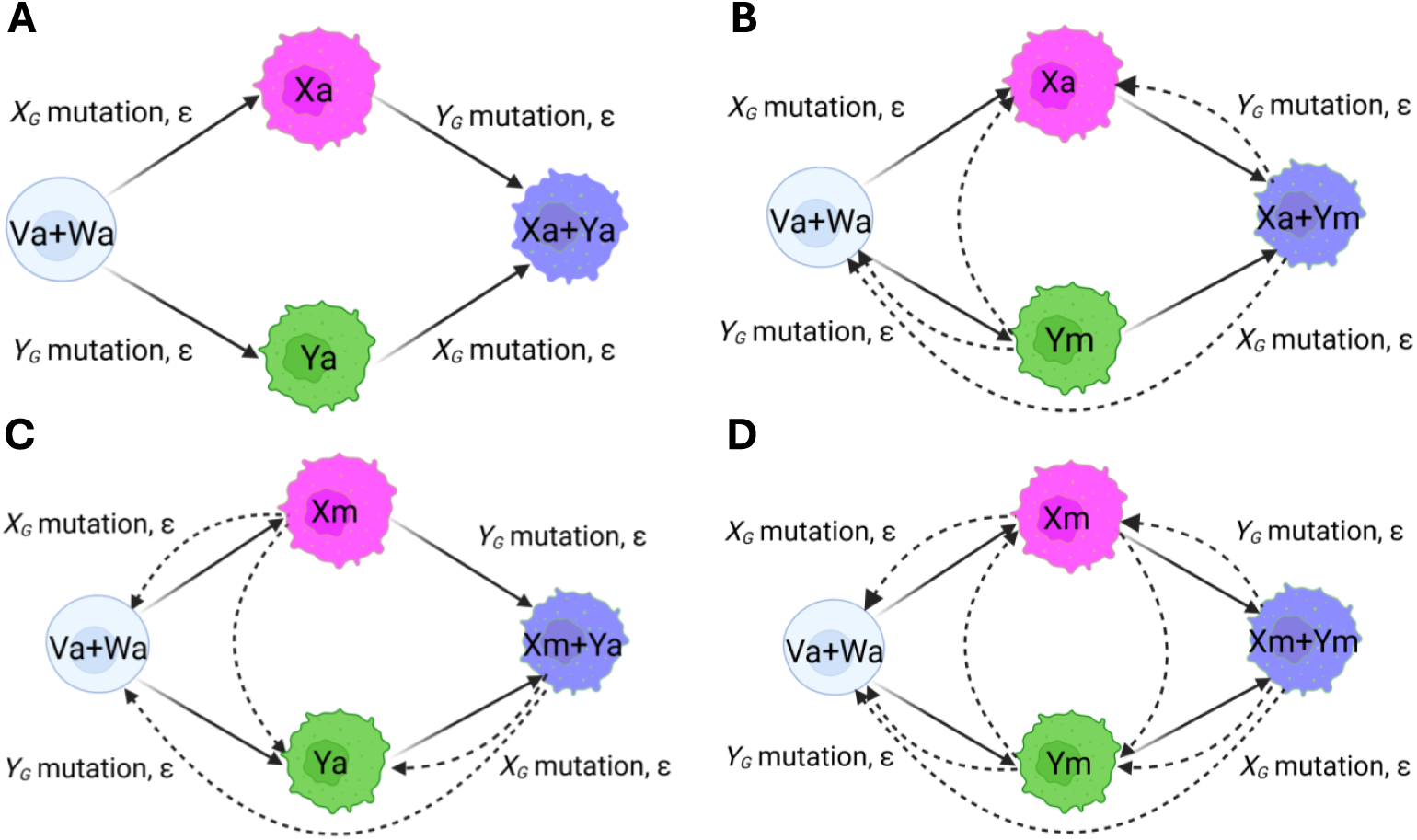
Schematic representation of the four evolutionary scenarios of epithelial carcinogenesis. In the illustration, healthy cells acquiring successive autonomous *V_G_* and *W_G_* mutations give rise to pre-malignant (*V_a_* + *W_a_*) cells. Solid line with arrow represents mutation, and dashed line with arrow represents stimulation of cells by the microenvironmental phenotype of a nearby cell. **A. Lineal evolution: Scenario XaYa** *V_a_* + *W_a_* cells either acquire autonomous *X_G_* or *Y_G_* mutations with probability *ε* and, consequently, give rise to *X_a_* or *Y_a_* cells, respectively. Then, these *X_a_* and *Y_a_* cells undergo subsequent autonomous mutations with the same probability, producing malignant (*X_a_* + *Y_a_*) cells. **B. Interclonal cooperativity: Scenario XaYm** *V_a_* + *W_a_* cells either acquire autonomous *X_G_* or microenvironmental *Y_G_* mutations with probability *ε* and thus, they give rise to *X_a_* or *Y_m_* cells, respectively. Afterwards, *X_a_* and *Y_m_* cells give rise to malignant *X_a_* + *Y_m_* cells by acquiring microenvironmental *Y_G_* and autonomous *X_G_* mutations, respectively, with the same probability. In this scenario, *X_a_* cells can be malignant as well, along with *X_a_* + *Y_m_* cells via the microenvironmental influence of *Y_m_* or *X_a_* + *Y_m_* cells. **C. Interclonal cooperativity: Scenario XmYa** *V_a_* + *W_a_* cells either acquire microenvironmental *X_G_* or autonomous *Y_G_* mutations with probability *ε* and produce *X_m_* or *Y_a_* cells, respectively. Then, with the same probability *ε*, *X_m_* and *Y_a_* cells undergo further autonomous *Y_G_* and microenvironmental *X_G_* mutations, respectively, and hence, produce malignant *X_m_* +*Y_a_* cells. In this scenario, *Y_a_* cells can be malignant as well via the microenvironmental influence of *X_m_* or *X_m_* + *Y_a_* cells. **D. Interclonal cooperativity: Scenario XmYm** The pre-malignant (*V_a_* + *W_a_*) cells either acquire microenvironmental *X_G_* or *Y_G_* mutations with probability *ε* and, thus, they give rise to *X_m_* or *Y_m_* cells, respectively. Then, with the same probability, *X_m_* and *Y_m_* cells acquire fourth microenvironmental *X_G_* and *Y_G_* mutations, respectively, and thus give rise to malignant *X_m_* + *Y_m_* cells. In this scenario, *V_a_* + *W_a_*, *X_m_*, and *Y_m_* cells can also be malignant via the microenvironmental influence of *X_m_*, *Y_m_* or *X_m_* + *Y_m_* cells. Figure created with BioRender.com (https://BioRender.com/z4m8vvc).

#### Malignancy of individual cells and lesions

We consider a cell to be malignant if it exhibits both hyperproliferative and invasive phenotypes, either by acquiring the mutations themselves or via microenvironmental effects from nearby cells [7, 8, 23]. Based on this, malignancy via lineal evolution can occur when a single cell clone acquires both mutations to become hyperproliferative and invasive. On the other hand, in malignancy by interclonal cooperativity, the required mutations do not need to occur within a single cell clone; rather, the mutations can occur in different cells, and they interact via their microenvironment to achieve malignancy. So, unless otherwise specified, the malignant cells produced in the four evolutionary scenarios (Figure 5) in our model are presented in Table 1.

**Table 1:**
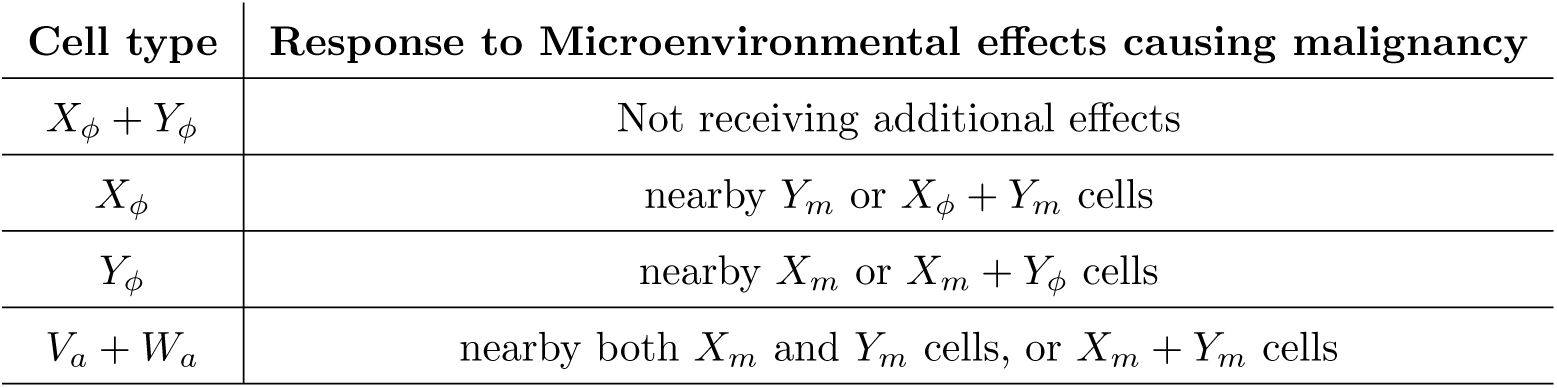
Different cell types that can be malignant under different microenvironmental conditions. Here, we define *ϕ ∈ {a, m}*, where *a* and *m* denote the cell phenotypes.

| **Cell type** | **Response to Microenvironmental effects causing malignancy** |
|---|---|
| *X*<sub>ϕ</sub> + *Y*<sub>ϕ</sub> | Not receiving additional effects |
| *X*<sub>ϕ</sub> | nearby *Y*<sub>*m*</sub> or *X*<sub>ϕ</sub> + *Y*<sub>*m*</sub> cells |
| *Y*<sub>ϕ</sub> | nearby *X*<sub>*m*</sub> or *X*<sub>*m*</sub> + *Y*<sub>ϕ</sub> cells |
| *V*<sub>*a*</sub> + *W*<sub>*a*</sub> | nearby both *X*<sub>*m*</sub> and *Y*<sub>*m*</sub> cells, or *X*<sub>*m*</sub> + *Y*<sub>*m*</sub> cells |

For an area of cells to be considered a “malignant lesion”, all of the cells in that area must satisfy 4 requirements simultaneously: (1) they are hyperproliferative, i.e. they have either an *X_a_* or *X_m_* phenotype, or they are within the range of influence of a nearby cell that has an *X_m_*genotype; (2) they are invasive, i.e. they have either a *Y_a_* or *Y_m_* phenotype, or they are within the range of influence of a nearby cell that has an *Y_m_* genotype; (3) they overlie an ECM that has been degraded at least to the critical threshold level, typically, 70%; (4) they are contiguous or connected and reach at least a certain threshold, typically, 100 cells.

In our model analysis, malignant lesion growth is quantified as the time required to reach the first target lesion size (typically 100 cells) under the above criterion. For simulations that do not reach the target within the observation time window, a value exceeding the maximum observation time is assigned for analysis.

#### Cross-talk interference

Cross-talk interference occurs when the activation of an intracellular signalling pathway by one type of mutation inhibits a signalling pathway activated by a second type of mutation in the same cell [7]. In the extreme case, one mutation completely inhibits the other, so there is no advantage of having the two mutations occurring in a single cell, and malignancy cannot be achieved through a lineal evolution trajectory. On the other hand, in the case of partial cross-talk interference, the *X_G_*and *Y_G_*mutations could occur in a single cell and still provide an incomplete proliferative or invasive advantage to the cell. To investigate varying levels of cross-talk interference, we devise our model such that if *ρ_V_* _+_*_W_* and *λ_V_* _+_*_W_* are the baseline proliferation and motility rates, respectively, of a *V* + *W* cell, then due to cross-talk interference, the cell with both *X_G_* and *Y_G_* mutations has a proliferation rate *ρ_X_*_+_*_Y_* = *ρ_V_* _+_*_W_* + (1 *− f*)*d_X_* and a motility rate *λ_X_*_+_*_Y_* = *λ_V_* _+_*_W_* + (1 *− f*)*m_Y_*, where *f* is an interference factor between 0 and 1, and *d_X_*and *m_Y_* are the increases in the proliferation and motility rate due to *X_G_*and *Y_G_*mutations alone, respectively, in that cell. In this case, *f* = 0 means no cross-talk interference, while *f* = 1 means extreme cross-talk interference.

#### Comprehensive model

Alongside the four 2-step scenarios shown in Figure 5, we built a comprehensive, multistep evolutionary model comprising all 16 cell types, possessing any combination of the traits *X_a_*, *X_m_*, *Y_a_*, and *Y_m_*. For example, a cell with none of the traits would be of type *V_a_*+*W_a_*, and a cell possessing all four traits would be of type *X_a_*+*X_m_*+*Y_a_*+*Y_m_*. We built this model with the same underlying rules for cell proliferation, movement, and death as before and with the same assumptions for two different phenotypic traits of mutant cells.

## 3 Results

### 3.1 Microenvironmental phenotypes accelerate carcinogenesis by enabling cooperation between nearby clones

To investigate the impact that the nature of the phenotype (cell autonomous or microenvironmental) has on carcinogenesis, we first used the VCBM to simulate carcinogenesis occurring through each of the four evolutionary scenarios shown in Figure 5 to determine which types of mutations–cell autonomous or microenvironmental–produce malignant lesions faster. We simulated Scenarios XaYa, XaYm, XmYa, and XmYm using the base parameter values in Table A1 and the modified values reflecting cell-cell interactions as specified in Tables B1– B4. From the simulations, we quantified both the number of cells of each genotype and the number of cells having a malignant phenotype, as defined in Section 2.1. We further quantified the percentage of the ECM area that had undergone at least 70% degradation. Representative outputs from the simulations are shown in Figure 6. The range and strength of interactions in *X_m_*and *Y_m_* cells were set at *r* = 60 µm (4-cell diameters) and *α* = 0.25, respectively.

**Figure 6:**
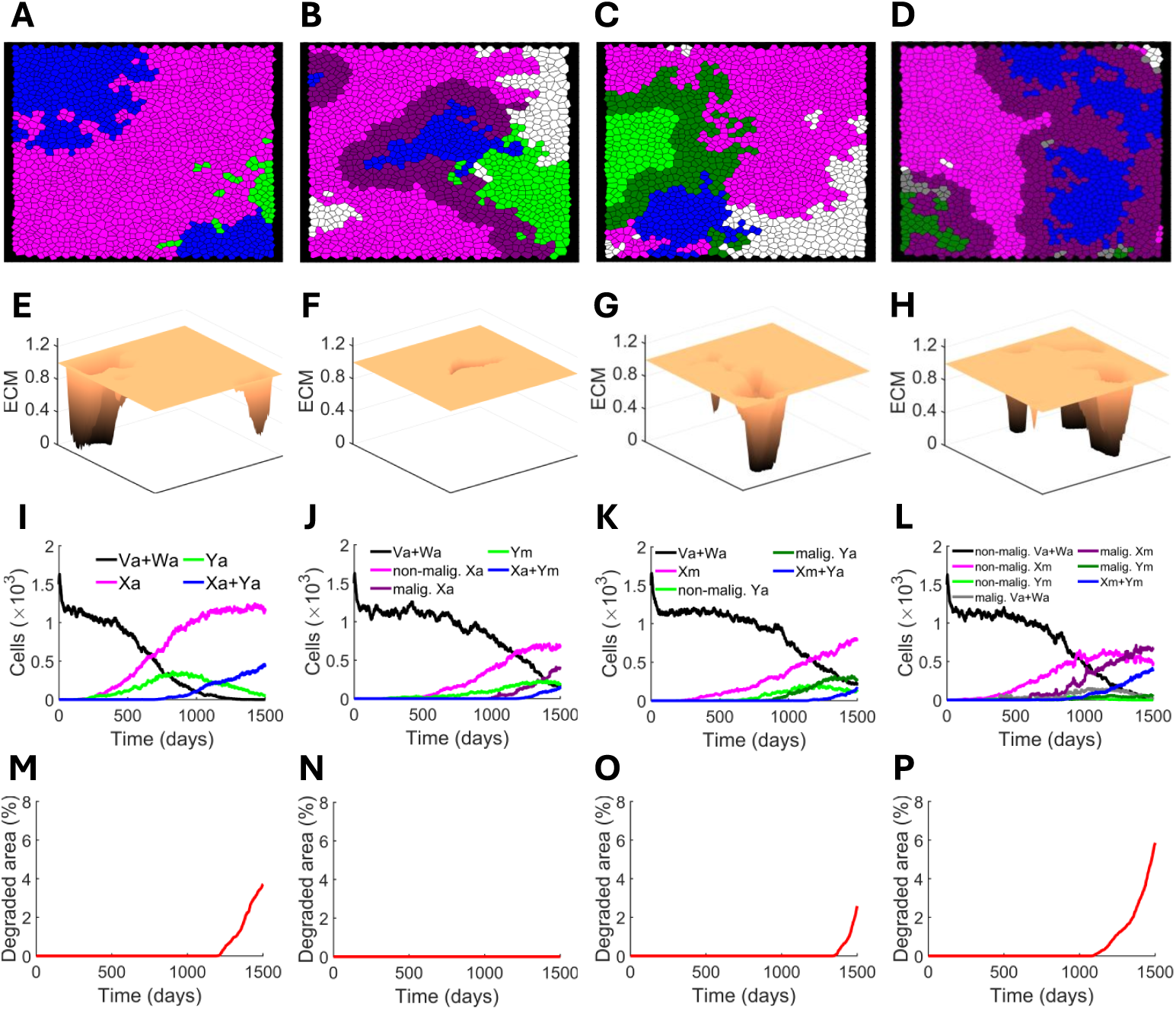
A VCBM of early carcinoma development. Outputs from a representative simulation of epithelial carcinogenesis evolution via Scenarios XaYa **(A, E, I, M)**, XaYm **(B, F, J, N)**, XmYa **(C, G, K, O)** and XmYm **(D, H, L, P)**. **A-D.** Voronoi tessellations at *t* = 1500 days. In the tessellations, (*V_a_* + *W_a_*) cells are white, cells with *X_G_* mutations are magenta, cells with *Y_G_* mutations are green and cells with both *X_G_* and *Y_G_* mutations are blue. In **B-D**, darker shades represent cells that are malignant under the influence of nearby cells harbouring *X_m_* and/or *Y_m_* mutations. **E-H.** The underlying ECM degradation profile at *t* = 1500 days. **I-L.** The time courses of the population of each cell type over 1500 days. The curves are colour-coded as in **A-D**, except for (*V_a_* + *W_a_*), which is shown in black. The legends shown in **I-L** apply to the corresponding panels **A-D**, respectively. **M-P.** The time courses of the area percentage with at least 70% ECM degradation, calculated as the percentage of locations with ECM values (*≤* 0.3) relative to the total number of locations in the domain at each time *t* for each of the four scenarios. The parameter values used for the simulations are those given in Tables A1 and B1-B4

We observed that the population of cells containing any *X_G_* or *Y_G_* mutation (i.e., the “mutant cell”) grew more rapidly in Scenario XaYa than in the other three scenarios, as seen as the mean number over time (Figure 7A) or measured as the inverse of the time taken to reach 100 cells (1*/t*_100_; Figure 7D). In contrast, the malignant cell population (which exhibits both hyperproliferative and invasive phenotypes) in Scenario XmYm grew significantly faster than in the other three scenarios (Figures 7B and E). Perhaps surprisingly, of the two scenarios that included a single microenvironmental phenotype, Scenario XaYm, which lacks a microenvironmental proliferation phenotype, produced malignant cells the fastest (Figure 7E). This highlights the importance of cell motility in the carcinogenic process. The underlying ECM of the carcinogenic lesion degraded over time across the four scenarios, as quantified by the percentage of the area exhibiting at least 70% ECM degradation. However, the ECM in Scenario XmYa degraded faster than in the other three scenarios (Figure 7C). We observed significant differences in malignant lesion-free survival among the four scenarios, as revealed by Kaplan-Meier analysis (log-rank test, *χ*^2^(3) = 74.86, *p <* 0.0001) (Figure 7F), where malignant lesion-free survival represents the percentage of simulations that had not yet developed a malignant lesion at a given time. However, Scenario XmYa showed an earlier decline in malignant lesion-free survival, decreasing from 100% to approximately 46% by 1490 days compared with other scenarios, suggesting earlier malignant lesion development in that scenario.

**Figure 7:**
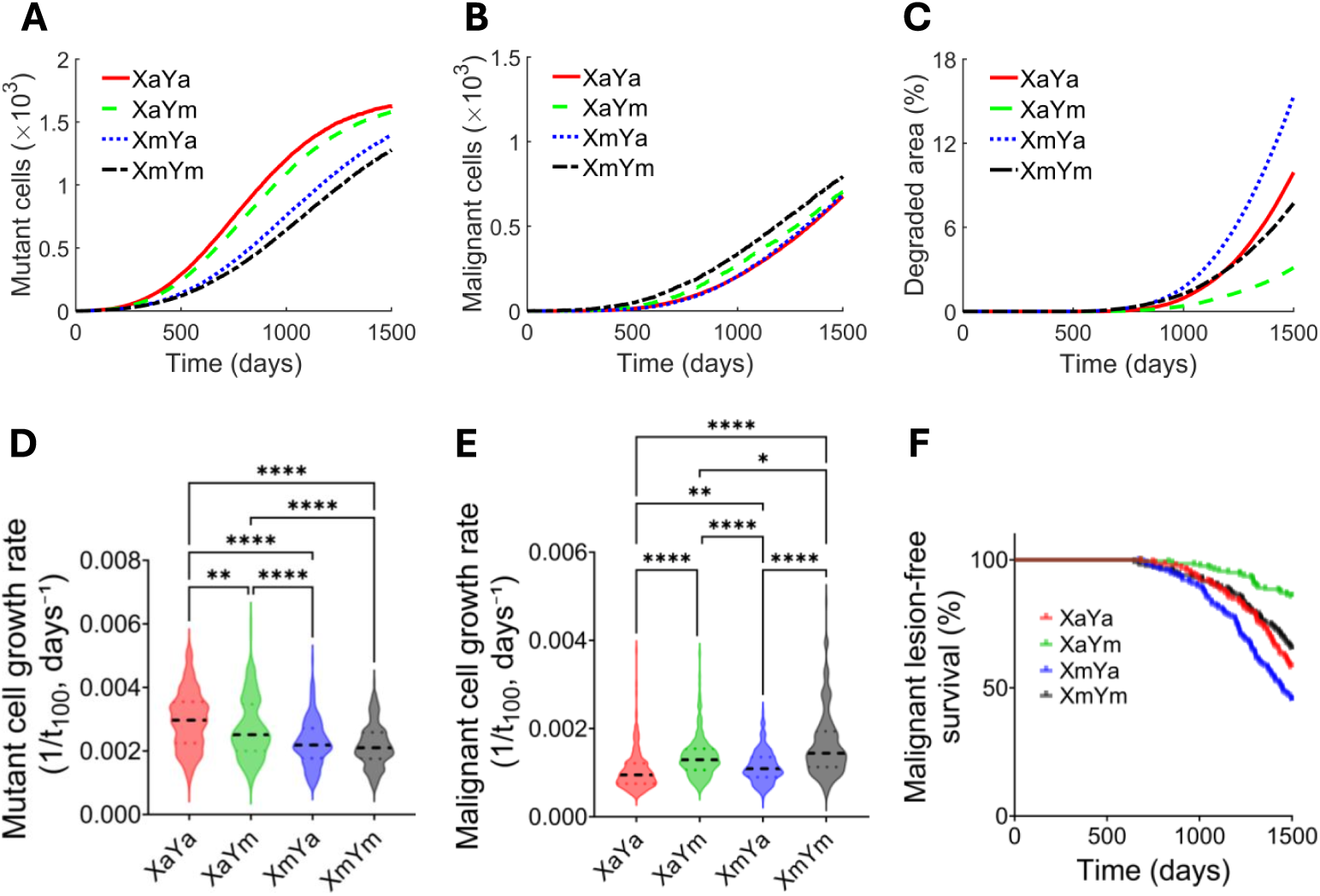
The impact of microenvironmental phenotypes on carcinogenesis. 200 carcinogenesis simulations were run for each of the four restricted evolutionary scenarios: XaYa, XaYm, XmYa and XmYm. The mean populations in each of the four scenarios over 1500 days are shown for mutant **(A)** and malignant **(B)** cells. **C.** The mean area of degraded ECM with at least 70% ECM degradation for each of the four scenarios is shown. **D-E.** The speed of growth of those simulations, measured as the inverse of the time to reach 100 mutant cells **(D)** or malignant cells **(E)**, is shown as violin plots with the median. **F.** Kaplan-Meier curves showing the percentage of simulations remaining free of malignant lesions (defined in Section 2.1) over time, with the development of the first malignant lesion considered the event of interest. Differences among the four scenarios were assessed using the log-rank test. Here and in all subsequent violin plots, the black dashed lines indicate the respective medians. The p-values for differences in **(D-E)** were determined using the Kruskal-Wallis test followed by Dunn’s multiple comparisons correction. Significant pairwise comparisons (*p ≤* 0.05) are indicated by asterisks.

We next investigated the effects of varying the range of the microenvironmental phenotypes, *r*, for a given strength of effect, *α* = 0.25, in Scenario XmYm. Interestingly, the growth rate of the mutant cell population decreased as the microenvironmental effect range increased by small amounts, but increasing the range beyond 4-cell diameters (i.e., 60 µm) had little further effect (Figures 8A and D). In contrast, the malignant cell population exhibited a modest increase in growth when both of the *X* and *Y* phenotypes were microenvironmental over a range of 2-cell diameters, but much more substantially as the range expanded to 4-, 6-, and 8-cell diameters (Figures 8B and E). Although overall mutant cell population growth decreased with the microenvironmental effect ranges, the underlying ECM degradation showed a non-monotonic dependence on this effect (Figure 8C). Malignant lesion-free survival differed significantly across the six varying ranges of the parameter *r* (Kaplan-Meier analysis, log-rank test, *χ*^2^(5) = 38.41, *p <* 0.0001), indicating sensitivity of the first malignant lesion development dynamics to this parameter (Figure 8F). However, the malignant lesion-free survival declined earlier in the range of *r* = 1-cell diameter, suggesting earlier malignant lesion development, which is consistent with the ECM degradation profile in Figure 8C.

**Figure 8:**
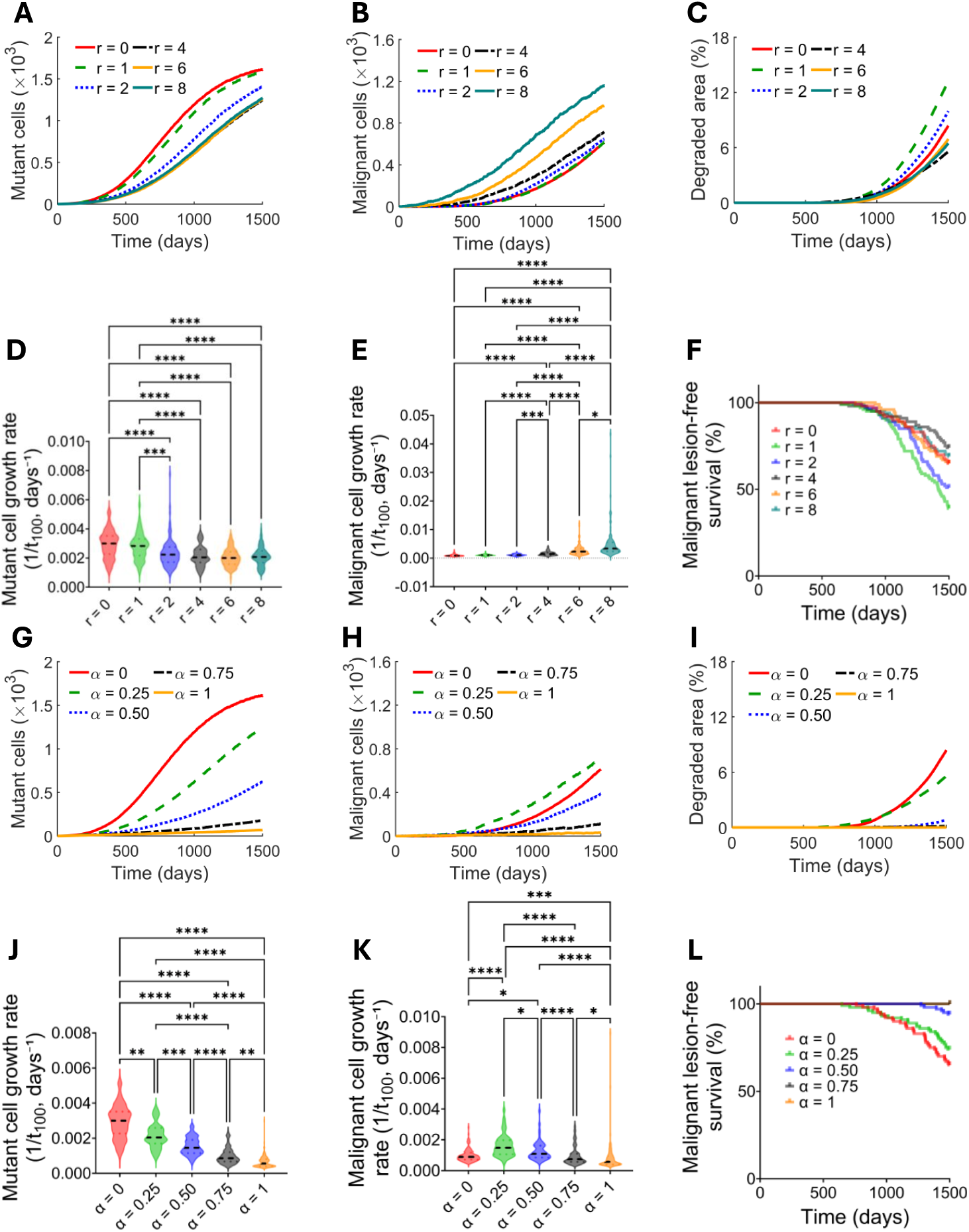
Clonal evolution in Scenario XmYm of epithelial carcinogenesis. The effects of varying the ranges **(A-F)** and strengths **(G-L)** of microenvironmental phenotypes on malignant characteristics are shown for Scenario XmYm. The mean number of mutant cells (**A** and **G**), number of malignant cells (**B** and **H**), and areas of degraded ECM with at least 70% ECM degradation (**C** and **I**) from over 100 simulations over time are shown. The speeds of acquiring malignant characteristics are represented by the inverse of the time elapsed to reach 100 mutant cells (**D** and **J**), 100 malignant cells (**E** and **K**) or by the time of first malignant lesion formation (**F** and **L**). The p-values for differences in **(D-E)** and **(J-K)** were determined using the Kruskal-Wallis test followed by Dunn’s multiple comparisons correction; significant differences (*p ≤* 0.05) are indicated by asterisks.

We then investigated the effects of varying the strength of the microenvironmental phenotypes, *α*, for a given range of effect, *r* = 4-cell diameters, in Scenario XmYm. The growth of mutant cell populations decreased dramatically as the strength of microenvironmental interactions increased (Figures 8G and J). In contrast, the malignant cell population increased significantly as the microenvironmental effect strength increased from 0 to 0.25, rising from a mean of 168 cells to 292 cells at *t* = 1000 days, where 0 means no interaction (Figures 8H and K). Surprisingly, as the strength of the microenvironmental effect increased to 0.50, the growth of the malignant cell population decreased dramatically, dropping from the mean of 292 cells to 123 cells at *t* = 1000 days. Increasing the strength to 0.75 and 1 further reduced the growth rate of the malignant cell population. The underlying ECM degradation decreased with increasing the strength of the microenvironmental effect, with the largest degradation observed in the absence of interaction at *α* = 0 (Figure 8I). This is consistent with the associated decrease in mutant cell population growth as the effect increases. The Kaplan-Meier analysis revealed significant differences in the malignant lesion-free survival across the five parameter values of *α* (log-rank test, *χ*^2^(4) = 94.87, *p <* 0.0001), with earlier decline observed at *α* = 0 (Figure 8L), consistent with the ECM degradation pattern in Figure 8I.

Similarly, we investigated the effects of varying the range and strength of the microenvironmental phenotypes, *r* and *α* (varying one at a time), on carcinogenesis using Scenarios XaYm and XmYa (Figures D1 and D2 in Appendix D). Scenarios XaYm and XmYa exhibited qualitative trends similar to those observed in Scenario XmYm for the analyses shown in Figures D1B, E, H and D2G, H, J, K. However, the remaining analyses displayed distinct behaviours.

Clonal cooperation can evolve through 3 basic types of cell lineage [7]: mother-daughter, in which a mutation is acquired by cells within a pre-malignant clone, giving rise to a new phenotype that complements that of the original cells; cousins, in which pre-malignant cells within a field independently acquire different mutations whose phenotypes are mutually beneficial; and neighbours, in which two clones located near each other in the tissue independently acquire mutations whose phenotypes are mutually beneficial. In the case of cousins and neighbours, the interacting clones begin as spatially separate fields of cells. To model this, we constructed two spatially separated patches of *X_m_* and *Y_m_* cells, each surrounded by (*V_a_* + *W_a_*) cells, and allowed them to grow in the absence of additional mutations. We observed that progression from premalignancy to malignancy can occur through field collision, i.e. by the growth and/or migration of mutually beneficial *X_m_* and *Y_m_* clones, leading to their co-localisation, which enables malignant interactions (Figure 9).

**Figure 9:**
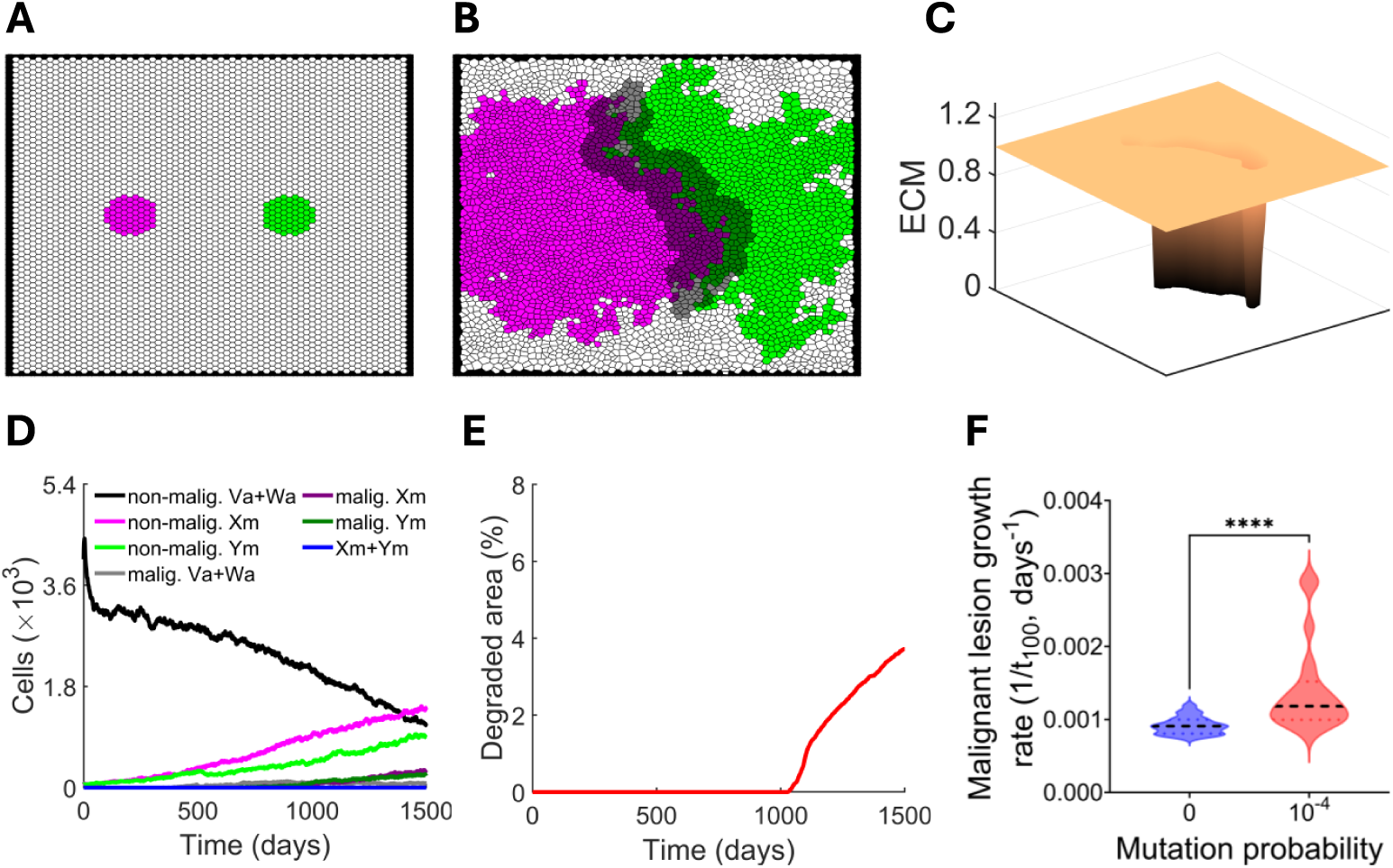
Progression to malignancy in the absence of mutation. **A.** To model carcinogenesis in the absence of mutagenesis, the tissue was seeded with 2 small fields of *X_m_* and *Y_m_* cells with interaction strength *α* = 0.25 and range *r* = 4-cell diameters, and a tissue-wide mutation rate, *ε* = 0. **B.** A representative image of the same tissue after 1500 days, by which time spatial expansion of the two initially separate cell fields collided. **C.** A representative image of the ECM underlying the tissue at 1500 days. **D.** The time courses of the population of each cell phenotype in the tissue over time. The curves are colour-coded as in **A-B**, except for (*V_a_* + *W_a_*), which is shown in black. The legends shown in **D** also apply to the panels **A** and **B**. **E.** The time course of the area percentage with at least 70% ECM degradation in the absence of mutation is shown. **F.** The speeds of malignant lesion formation for the standard (10*^−^*^4^) probability of acquiring a mutation and the absence (0) of a mutation. Data in **F** are from 20 simulations. The p-value for the difference was determined using the Mann-Whitney U-test (*U* = 49, *p <* 0.0001, 20 observations), with statistical significance indicated by asterisks.

### 3.2 Clonal evolution in a comprehensive genetic model of epithelial carcinogenesis

Next, we simulated epithelial carcinogenesis using the comprehensive model, in which all four mutations that cause cell-autonomous or microenvironmental phenotypes for growth (*X_a_*and *X_m_*) and invasion (*Y_a_*and *Y_m_*) could be acquired with equal probability and in any order. A representative endpoint diagram, an evolution plot and the underlying ECM degradation are shown in Figure 10. In these simulations, when either the strength of interaction, *α*, or its range, *r*, was set to zero, only cell-autonomous phenotypes were present, i.e., *X_m_* and *X_a_* were equivalent, as were *Y_m_*and *Y_a_*.

**Figure 10:**
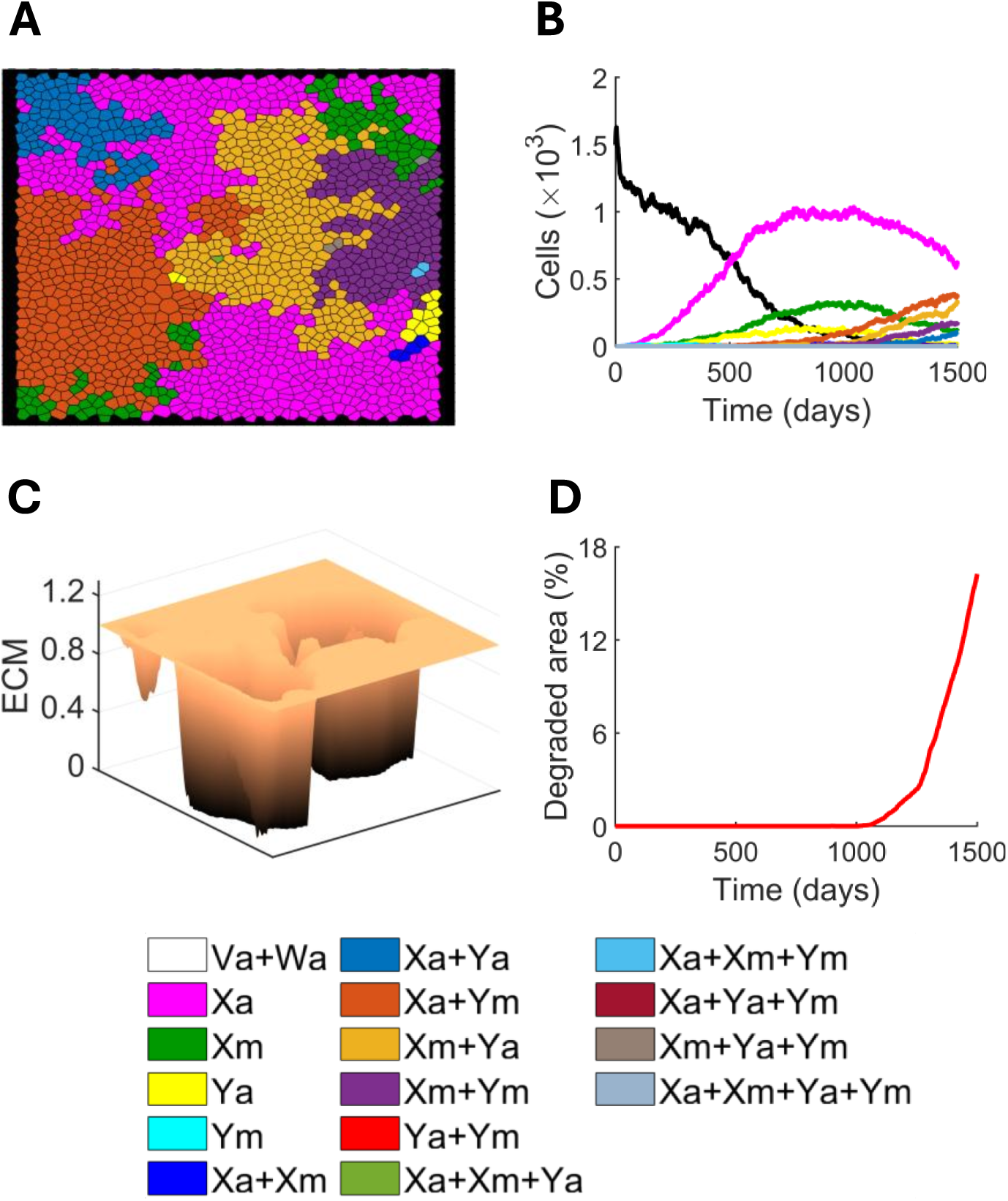
Outputs from a representative simulation of epithelial carcinogenesis evolution via the Comprehensive model **(A, B, C, D)**. **A.** Voronoi tessellation at *t* = 1500 days. **B.** The time courses of the population of each cell type over 1500 days. **C.** The underlying ECM degradation profile at *t* = 1500 days. The curves are colour-coded as in **A**, except for *V_a_* + *W_a_*, which is shown in black. **D.** The time course of the percentage of area with at least 70% ECM degradation. The parameter values used for the simulations are those given in Tables B1-B4 (for ECM degradation only) and B5.

We began by investigating the effect of varying the range of the microenvironmental phenotypes, *r*, at a given strength of effect, *α* = 0.25, in the comprehensive model. The growth rate of the mutant cell population decreased slightly as the microenvironmental effect range increased from 0 to 1-, 2-, and 4-cell diameters, but further increases beyond 4-cell diameters had only a minimal effect (Figures 11A and D). In contrast, the malignant cell population increased as the microenvironmental effect was influenced over a range of 2-cell diameters, rising from approximately 404 cells to 520 cells at *t* = 1000 days, with only minor further increases up to 6-cell diameters and a more substantial increase at 8-cell diameters (Figures 11B and E). On the other hand, the ECM degradation exhibited distinct temporal profiles across the microenvironmental effect ranges, with only slight differences between 1- and 2-cell diameters, lower degradation at 4- and 6-cell diameters, and intermediate degradation at 8-cell diameters (Figure 11C). Malignant lesion-free survival differed significantly for the six varying ranges of *r* (Kaplan-Meier analysis, log-rank test, *χ*^2^(5) = 34.04, *p <* 0.0001), with malignant lesion-free survival declining slightly earlier at the range of *r* = 1-cell diameter, suggesting slightly earlier malignant lesion development (Figure 11F), consistent with the ECM degradation pattern in Figure 11C.

**Figure 11:**
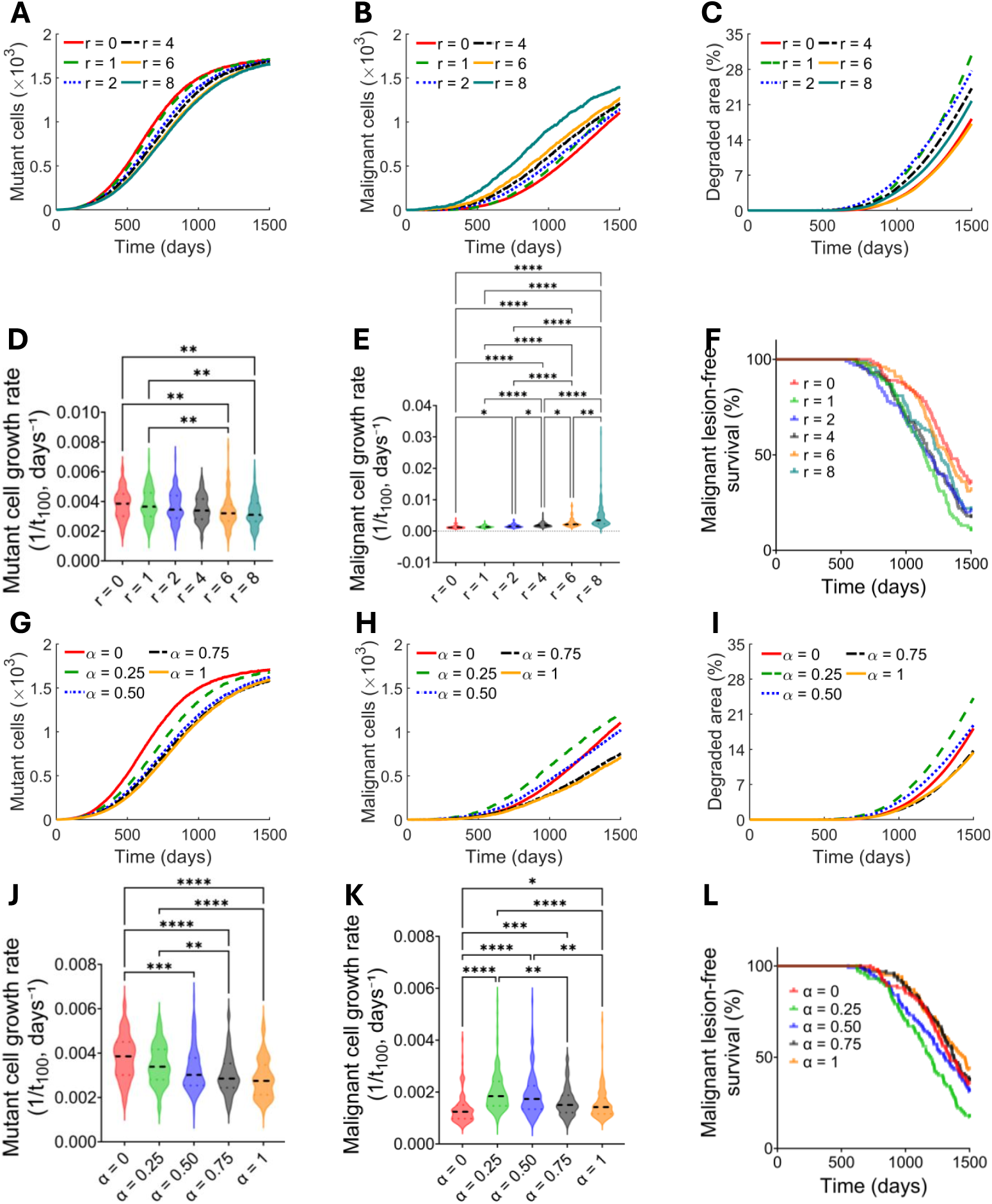
Clonal evolution in the comprehensive genetic model of epithelial carcinogenesis. The effects of varying the ranges **(A-F)** and strengths **(G-N)** of microenvironmental phenotypes on malignant characteristics in the comprehensive evolutionary model. The mean number of mutant cells (**A** and **G**), number of malignant cells (**B** and **H**) and areas of degraded ECM with at least 70% ECM degradation (**C** and **I**) from over 100 simulations over time are shown. The speeds of acquiring malignant characteristics are represented by the inverse of the time elapsed to reach 100 mutant cells (**D** and **J**), 100 malignant cells (**E** and **K**) or by the time of first malignant lesion formation (**F** and **L**). The *p*-values for differences in **(D-E)** and **(J-K)** were determined using the Kruskal-Wallis test followed by Dunn’s multiple comparisons correction; significant differences (*p ≤* 0.05) are indicated by asterisks.

We next investigated the effect of varying the strength of microenvironmental interaction, *α*, in the comprehensive model, while keeping a fixed range of effect (*r* = 4-cell diameters). As *α* increased from 0 to 0.25 and 0.50, we found that the growth of the mutant cell population slowed markedly, whereas further increases in *α* to 0.75 and 1 resulted in a similar reduction in growth (Figures 11G and J). In contrast, the growth rate of the malignant cell population increased when the microenvironmental effect of strength increased from 0 to 0.25, rising from a mean of 404 cells to 604 cells at *t* = 1000 days (Figures 11H and K) but then decreasing dramatically as the effect strength increased to 0.50, 0.75 and 1, with only slight differences in the temporal profiles for 0.75 and 1, dropping from approximately 604 to 452, 288 and 283 cells, respectively, at *t* = 1000 days. The effects of varying *r* and *α* on mutant and malignant cell population growth were qualitatively consistent between Scenario XmYm and the comprehensive model, with only differences in the magnitude of the response. On the other hand, the ECM degradation increased markedly as the microenviron-mental effect strength increased from 0 to 0.25, then decreased with further increases in the effect strength, with strengths 0.75 and 1 exhibiting similar levels of degradation (Figure 11I). The Kaplan-Meier analysis revealed significant differences in the malignant lesion-free survival across the five values of *α* (log-rank test, *χ*^2^(4) = 27.70, *p <* 0.0001), with malignant lesion-free survival declining earlier at *α* = 0.25, decreasing from 100% to 17% by 1498 days (Figure 11L), suggesting earlier malignant lesion development. This is consistent with the ECM degradation pattern in Figure 11I.

We then quantified the composition of cell types in the initial malignant lesions (defined in Section 2.1) across 126 simulations. The frequency with which each cell type formed the majority of the malignant lesions was quantified across simulations. The cell type *X_a_* + *Y_a_*was most frequently dominant, occurring in approximately 30% of simulations followed by *X_a_* +*Y_m_* (approximately 21%); *X_m_* +*Y_a_* cell type dominated in approximately 17% of simulations, *X_m_* + *Y_m_*cell type was dominant in approximately 9%, while *Y_a_* and *Y_m_* cell types did so in approximately 13% and 10%, respectively (Figure 12B). The remaining cell types did not achieve dominance. In contrast, multiple cell types were commonly present within malignant lesions, with *X_a_* cell type being the most frequent, appearing in approximately 56% of simulations (Figure 12A). To characterise the diversity of malignant lesions, we quantified the frequency of pairwise co-occurrence among the intrinsic phenotypes (Figure 12C). The heat map revealed distinct patterns of pairwise co-occurrence, with some phenotype combinations occurring more frequently than others. In particular, combinations involving phenotypes *V_a_* + *W_a_*and *X_a_*were among the most frequently observed, whereas those involving phenotype *X_m_* + *Y_m_*generally occurred at lower frequencies.

**Figure 12:**
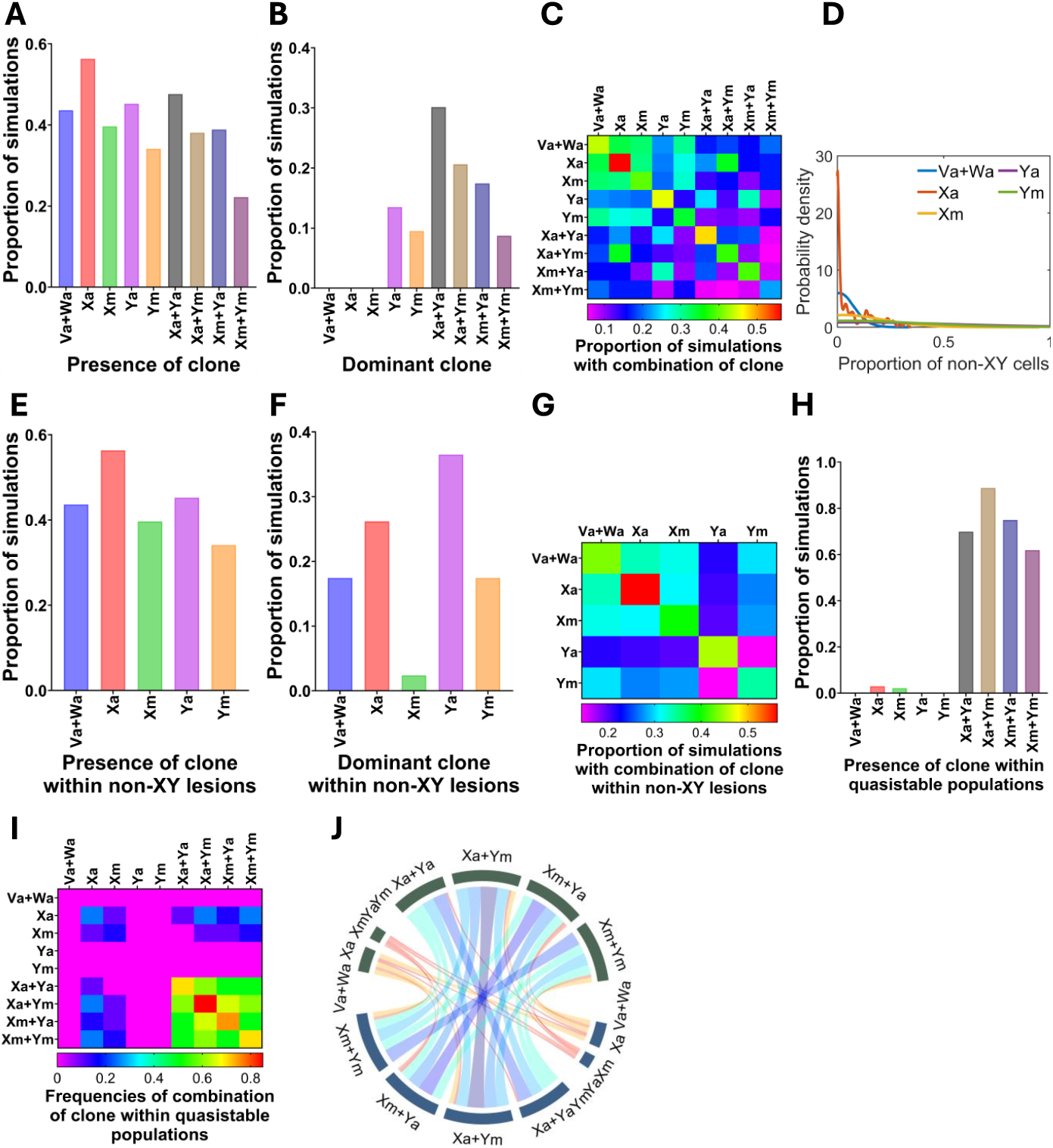
Genetic and phenotypic heterogeneity within the malignant lesions arising in the comprehensive model. Frequencies and composition of nine intrinsic phenotypes (genotypes with the same intrinsic phenotype were pooled) in the initial malignant lesions across 126 simulations. This analysis is conducted with interaction strength *α* = 0.25 and range *r* = 4-cell diameters. **A.** Proportion of simulations in which each clone intrinsic phenotype is present within malignant lesions. **B.** Proportion of simulations in which each clone phenotype within malignant lesions is dominant. Dominance is defined as having the highest proportion in a given simulation. **C.** Heat-map showing relative proportions of simulations in which 2-way combinations of intrinsic phenotypes co-exist in the malignant lesions. **D.** Kernel density estimation plot of the proportion of non-XY cells across simulations. **E.** Proportion of simulations in which each non-XY clone intrinsic phenotype is present within malignant lesions. **F.** Proportion of simulations in which each non-XY clone intrinsic phenotype is dominant. **G.** Heat-map showing relative proportions of simulations in which 2-way combinations of non-XY clone phenotypes co-exist in the lesions. **H.** Proportion of simulations in which each clone intrinsic phenotype is present within the quasistable malignant lesions at a minimum of 5% of the cell population. **I.** Heat-map showing relative proportions of simulations in which 2-way combinations of intrinsic phenotypes co-exist in the quasistable malignant populations. **J.** Chord diagram representation of each intrinsic phenotype and the others that are found

To characterise the distribution of non-XY intrinsic phenotype proportions in malignant lesions, we constructed kernel density estimates, which showed that the distributions varied among intrinsic phenotypes across simulations (Figure 12D). Phenotype *X_a_* was strongly concentrated near zero, indicating that this phenotype was absent or present at very low proportions in most simulations. In contrast, phenotype *V_a_* + *W_a_*exhibited a broader distribution. The remaining non-XY phenotypes were generally concentrated at low proportions with high right tails, indicating that high proportions occurred in relatively few simulations. The non-XY intrinsic phenotypes contributed to malignant lesions at different levels (Figure 12E), with the *Y_a_* phenotype predominating and appearing as dominant in approximately 37% of simulations, followed by *X_a_*(26%), while *V_a_* + *W_a_* and *Y_a_* were each dominant in approximately 17% of simulations (Figure 12F). This indicates that the greater invasiveness capacity of *Y_a_* cells confers a competitive advantage, enabling them to dominate malignant lesions. The heat map revealed that combinations involving phenotypes *V_a_* + *W_a_* and *X_a_* were among the most frequently observed, while those involving phenotype *Y_a_* occurred less frequently among non-XY intrinsic phenotypes in the initial malignant lesions (Figure 12G).

In simulations that were allowed to continue beyond 1500 days, it was noticed that, after a period of rapid change and clonal succession, groups of cell types persisted together in the tissue at similar levels for long periods of time, leading to the entire tissue being malignant (Figure C5P). We quantified the proportions of each cell genotype with a proportion of at least 5% in tissues in this quasistable state over *t* = 2500-3500 days across 20 simulations. The four intrinsic phenotypes, including *X_a_* +*Y_a_*, *X_a_* +*Y_m_*, *X_m_* +*Y_a_* and *X_m_* +*Y_m_*, were most frequently observed, appearing in approximately 62-89% of the quasistable phase (Figure 12H). Cell types *X_a_* and *X_m_* were rarely present (less than 5%), suggesting minimal contribution. The heat map revealed that combinations involving four intrinsic phenotypes: *X_a_* + *Y_a_*, *X_a_* + *Y_m_*, *X_m_* + *Y_a_*and *X_m_* + *Y_m_*were among the most frequently observed, while combinations involving phenotypes *X_a_* and *X_a_* occurred less frequently (Figure 12I). Consistent with this pattern, the chord diagram showed strong connections among phenotypes *X_a_* +*Y_a_*, *X_a_* + *Y_m_*, *X_m_* + *Y_a_* and *X_m_* + *Y_m_*, indicating frequent pairwise co-occurrence in the quasistable phase (Figure 12J).

### 3.3 Signalling pathway interference

Many oncogenic mutations drive carcinogenesis by dysregulating intracellular signal transduction pathways that affect the proliferative and invasive behaviours of the cells. However, the activation of some pathways can interfere with the activity of other pro-malignant pathways [58–61]. Therefore, we investigated the effect on cancer cell evolution of varying levels of pathway cross-talk interference in the comprehensive model, where the enhancement of proliferation by *X_a_* is multiplied by the interference factor (1 *− f*), where 0 *≤ f ≤* 1, when the cell also has a *Y_a_*mutation. Similarly, the enhancement of motility by *Y_a_* is multiplied by (1 *− f*) when the cell also has an *X_a_*mutation. Thus, when *f* = 0, there is no pathway interference between *X_a_* and *Y_a_*, and when *f* = 1, there is complete interference.

The number of mutant cells changed little with the degree of interference (Figures 13A and D). In contrast, a 25% level of interference (*f* = 0.25) caused a dramatic reduction in the number of malignant cells, which was further reduced as interference levels rose, with a similar reduction in growth observed for higher interferences (*f* = 0.75 and *f* = 1) (Figures 13B and E). Similarly, the ECM degradation decreased progressively as interference increased (Figure 13C). The malignant lesion-free survival differed significantly across varying interference levels *f* (Kaplan-Meier analysis, log-rank test, *χ*^2^(4) = 212.7, *p <* 0.0001); however, the malignant lesion-free survival declined significantly earlier at *f* = 0, followed by *f* = 0.25 and *f* = 0.50, suggesting earlier malignant lesion development (Figure 13F). On the other hand, simulations with *f* = 0.75 and 1 did not reach the threshold for a malignant lesion within 1500 days over 50 simulations. As a result, at those values, the malignant lesion-free survival was always 100% over time.

**Figure 13:**
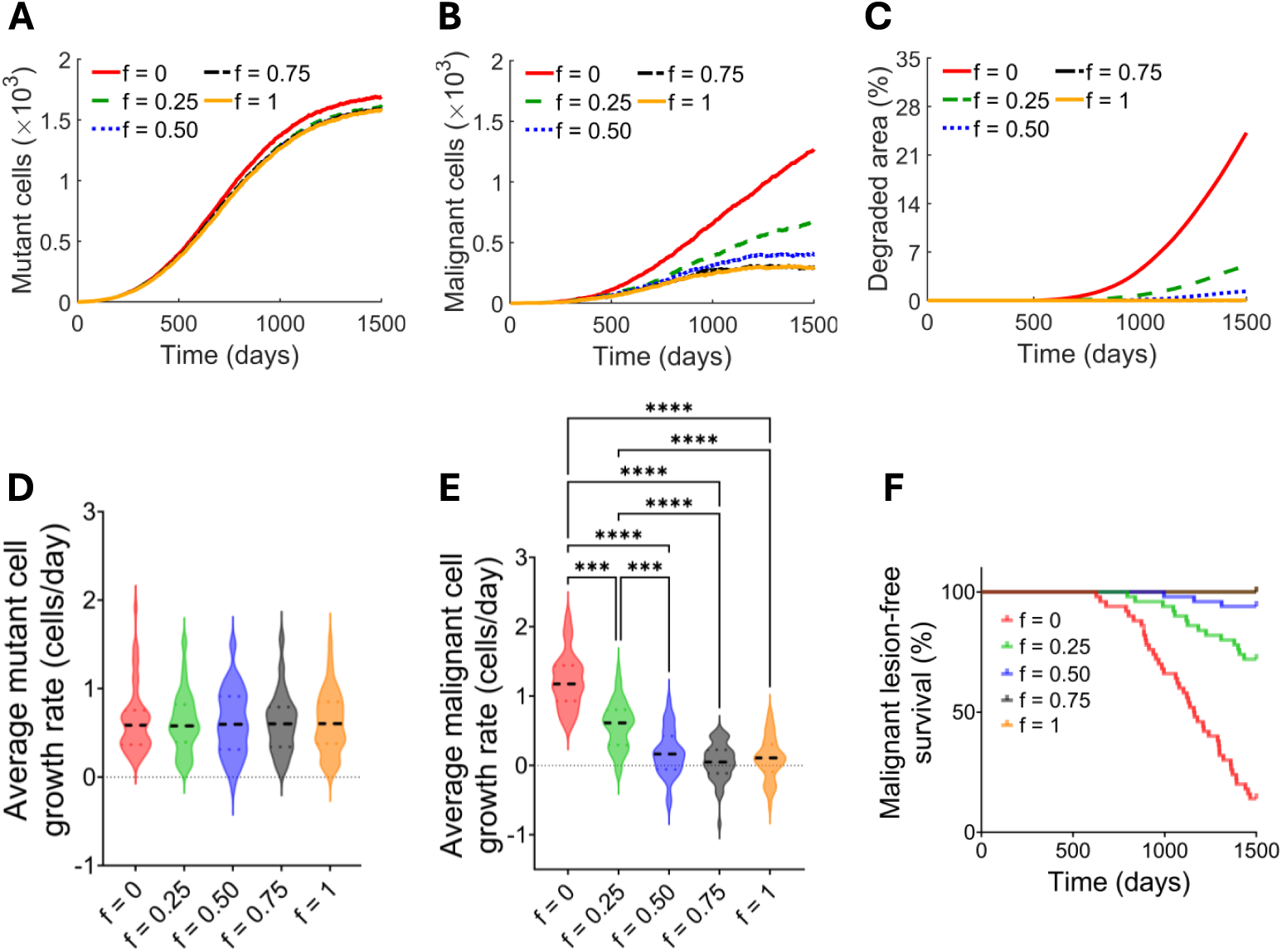
Effects of intracellular signalling pathway interference on carcinogenesis. Mean mutant **(A)** and malignant **(B)** cell populations are shown for different levels of pathway interference in the comprehensive model, keeping *α* constant at 0.25, and the interaction range *r* constant at 4-cell diameters. **C.** The mean area of degraded ECM with at least 70% ECM degradation for various levels of interference. Average growth rates of mutant cells **(D)** and malignant cells **(E)** between 1000 and 1500 days, calculated as ((*N*_1500_ *−N*_1000_)*/*(1500*−*1000)), where (*N_t_*) denotes the number of cells at time *t*, are shown as violin plots with the median (black dashed). **F.** The Kaplan-Meier curves of malignant lesion-free survival for different levels of interference. Results are averaged over 50 simulations across 1500 days. The p-values for differences in **D** and **E** were determined using the Kruskal-Wallis test followed by Dunn’s multiple comparisons correction. Only pairwise comparisons with *p ≤* 0.05 are shown by asterisks.

## 4 Discussion

Early tumour detection can lead to complete eradication or prevention of cancer. Lack of a proper understanding of tumour mechanisms, such as tumour formation via a succession of mutations, and heterogeneous cell and tissue characteristics in tumour progression, is the main barrier to implementing an efficient treatment strategy. In this study, we developed a VCBM for the interactions and cooperation between hyperproliferative and invasive clones (produced by mutations) and investigated both evolutionary paradigms (lineal evolution and interclonal cooperativity) in driving premalignant tissue to malignant lesions that progress to cancer. Our results indicate that mutations producing microenvironmentally extended phenotypes in interclonal cooperativity trajectories can accelerate malignancy compared with cell-autonomous phenotypes in the lineal evolution trajectory, through symbiotic clonal interactions.

Cells can produce malignant lesions via lineal evolution or interclonal cooperativity scenarios while invading the underlying ECM. We simulated these evolutionary scenarios (i.e., four scenarios) of carcinogenic lesion progression, as well as the comprehensive genetic model comprising all 16 cell types, each possessing any combination of the traits *X_a_*, *X_m_*, *Y_a_*, and *Y_m_*.

Our results indicate that Scenario XaYa favours early mutant cell expansion (Figures 7A and D). This behaviour may arise because the more abundant pre-malignant cells in the three scenarios with microenvironmentally extended phenotypes (i.e., scenarios XaYm, XmYa, and XmYm) are more likely to benefit from microenvironmental interactions, enabling their prolonged persistence compared to those in Scenario XaYa (see Figures 6A-D). Consequently, mutant cell growth is reduced in these scenarios relative to Scenario XaYa. In contrast, Scenario XmYm promotes early malignant cell progression (Figures 7B and E). This can be attributed to the presence of both *X_G_* and *Y_G_* mutations in Scenario XmYm, which give rise to microenvironmental phenotypes; cells carrying these mutations may affect many cells in their vicinity by increasing their proliferation and motility. Together, these findings highlight the role of cellular cooperation on tumour dynamics and are consistent with the experimental observations reported by Guzelsoy *et al.* [62]. Surprisingly, among scenarios with a single microenvironmental phenotype, Scenario XaYm, which lacks a microenvironmental proliferation phenotype, produced malignant cells most rapidly, highlighting the importance of cell motility in carcinogenesis. This finding is consistent with those reported by Gerlee and Anderson [63]. The underlying ECM degraded over time across the four scenarios; however, the ECM in Scenario XmYa degraded faster than in the other three scenarios (Figure 7C). Consistent with this, malignant lesion development occurs slightly earlier in Scenario XmYa than in the other scenarios (Figure 7F). The reason may be that ECM degradation is a key criterion for malignant lesion formation (as defined in Section 2.1), and occurs at an increased rate in scenarios XmYa and XmYm due to the microenvironmental influence of hyperproliferative cells (i.e., *X_m_* cells). Therefore, we would expect faster ECM degradation and, consequently, earlier malignant lesion development in Scenario XmYm. However, this expectation does not account for the fact that the pre-malignant cells that become malignant in Scenario XmYm do not contribute to ECM degradation, which may help explain the observed pattern of malignant lesion development. Overall, these findings suggest that the acquisition of hyperproliferative capacity may confer a modest growth advantage during the early stages of tumour progression by accelerating ECM degradation through microenvironmental interactions.

We next investigated cell population growth as a function of the microenvironmental effect range and strength parameters, *r* and *α*, respectively. Our results indicate that as *r* increases up to 4-cell diameters, the mutant cell population growth decreases slightly; however, further increases in the interaction range have little additional effect on growth, suggesting that the influence of the microenvironmental effect range reaches a plateau beyond this threshold (Figures 8A and D, and 11A and D). Likewise, as *α* increases, the mutant cell population growth markedly decreases in both Scenario XmYm and the comprehensive model (Figures 8G and J, and 11G and J); however, the reduction becomes minimal in the comprehensive model at an effect strength higher than 0.5. One possible explanation for these findings is that more abundant pre-malignant cells may benefit more from microenvironmental interactions than mutant cells as the interaction range and strength of effect increase. In contrast, the malignant cell population growth increases modestly as *r* increases from 0 to 4-cell diameters and beyond (Figures 8B and E, and 11B and E). The malignant cell growth also increases at lower values of the interaction strength (*α*), but decreases when the strength of the effect becomes sufficiently large, e.g., at 0.5 and higher (Figures 8H and K, and 11H and K). This behaviour may arise because, as the strength of microenvironmental interactions in a cell increases, the most abundant pre-malignant cells (*V_a_* + *W_a_*) within *r* = 4-cell diameters of that particular cell move and proliferate more efficiently, ultimately outcompeting and crowding out the microenvironmental stimulator cell. This reduces mutant cell population growth and, consequently, that of malignant cells (see Figures D3A-L in Appendix C). Overall, this pattern in malignant cell population growth can be explained by the fact that a larger interaction range allows more cells to respond to microenvironmental effects, thereby promoting more malignant cell growth. On the other hand, a moderate increase in interaction strength enhances this growth. Still, at higher values, it strengthens competition from the increased abundance of surrounding pre-malignant cells, which ultimately limits malignant cell expansion. However, further investigation is required to confirm these hypotheses. In ECM degradation, Scenario XmYm exhibits a non-monotonic dependence on the microenvironmental effect range (*r*), but a monotonic decrease with the effect strength (*α*). Similarly, the comprehensive model shows a non-monotonic dependence on *r* in ECM degradation, but a monotonic decrease at 0.5 and higher, with minimal changes at 0.75 and 1 (Figures 8C and I, and 11C and I). The Kaplan-Meier analysis reveals significant differences in the timing of first malignant lesion development across different microenvironmental interaction ranges and strengths in both Scenario XmYm and the comprehensive model. The malignant lesion development patterns across different ranges and strengths are consistent with ECM degradation, a key criterion for malignant lesion formation.

In our model, we considered a scenario with two spatially separated patches of *X_m_* and *Y_m_* cells, each surrounded by *V_a_* +*W_a_* cells, in the absence of additional mutations. The results suggest that progression from pre-malignancy to malignancy can occur via field collision, whereby the growth and/or migration of mutually beneficial *X_m_* and *Y_m_*clones leads to their co-localisation, enabling malignant interactions (Figure 9). This highlights field collision as a potential pathway to malignancy even in the absence of additional mutations and emphasises the role of spatial structure and clonal co-operation via microenvironmentally extended phenotypes in driving malignant lesion progression.

We further assessed the frequencies and composition of cell types in the initial malignant lesions at time *t* = 800 days across 126 simulations using the comprehensive model. The findings indicate that the greater invasiveness capacity of *Y_a_*cells confers a competitive advantage, enabling them to dominate initial malignant lesions among non-XY phenotypes in approximately 37% of simulations (Figure 12F). The findings also show that combinations involving the phenotypes *V_a_* + *W_a_*and *X_a_*were among the most frequently observed pairwise co-occurrences among the intrinsic phenotypes, indicating their frequent association during malignant lesion progression (Figures 12C and G).

Following a period of rapid change and clonal succession, the system converges to a quasistable state characterised by the long-term coexistence of multiple cell types and the entire tissue being malignant. Analysis over *t* = 2500–3500 days across 20 simulations indicates that four intrinsic phenotypes *X_a_* + *Y_a_*, *X_a_* + *Y_m_*, *X_m_* + *Y_a_*and *X_m_* + *Y_m_*, were most frequently observed, contributing in a mean of approximately 62-89% of the quasistable phase (Figure 12H), whereas phenotypes (*X_a_*and *X_m_*) were observed only infrequently and other were not observed, suggesting a limited role in maintaining quasistability.

Finally, using the comprehensive model, we investigated the effect of cross-talk interference in the cell signalling pathway, defined by *f*. The results indicate limited change in mutant cell growth with *f*, indicating a minimal effect of interference on the overall mutant cell growth. In contrast, malignant cell abundance, malignant lesion development, and ECM degradation decreased gradually with *f*, consistent with higher values corresponding to strong cross-talk interference, hence reducing lesion growth and matrix degradation. At higher *f*, malignant cells were produced primarily through microenvironmental influences. This suggests that microenvironmental phenotypes may overcome the limits on carcinogenesis caused by mutual inhibition between pro-malignant intracellular signalling pathways.

As we build the VCBM of epithelial carcinogenesis evolution and explore a tissue that is just becoming carcinogenic in this study, we plan to extend this model to more advanced stages of carcinogenesis and treatment strategies in the future. Due to computational costs, we excluded broad investigation and measurement of the invasion of the underlying ECM by cells in the current work; therefore, we plan to conduct such studies in future work. Cell heterogeneity, arising from mutations, diverse cellular characteristics and behaviours, cell-cell cooperation, and cell-ECM dynamics that drive the progression of malignant lesions to cancer, has become an increasingly important focus of research. The present study offers a framework for realistically modelling all these factors in the earliest stages of tumour progression.

## 5 Conclusion

Cancer progression from a succession of mutations is an evolutionary process that takes a highly variable time to produce a carcinogenic lesion that progresses to cancer. The evolutionary process can be of two types: lineal evolution, in which mutations produce non-cooperative cell-autonomous phenotypes, or interclonal cooperativity, in which mutations produce cooperative microenvironmentally extended phenotypes. The *in silico* model developed in this study shows how carcinogenic lesions form along these evolutionary processes and indicates which of these can lead to faster malignant lesion progression by extensively degrading the underlying ECM. This study demonstrates that mutations producing microenvironmentally extended phenotypes via interclonal cooperativity can accelerate the onset of malignancy compared with cell-autonomous phenotypes via lineal evolution, through symbiotic clonal interactions. Furthermore, progression from pre-malignancy to malignancy can occur without the acquisition of additional mutations, through field collision, i.e. by the growth and/or migration of mutually beneficial clones, leading to their co-localisation, which enables malignant interactions. In addition, following a period of rapid change and clonal succession, groups of cell types persist together for long periods, maintaining stable proportions. Finally, microenvironmental phenotypes can overcome the limits on carcinogenesis caused by mutual inhibition of pro-malignant intracellular signalling pathways. Together, these findings highlight the critical role of microenvironmental interactions in cancer evolution. The computational model developed in this study provides a useful tool for investigating the role of cell cooperation in malignant lesion growth during the earliest stages of carcinogenesis and may support future biological and clinical research across different cancer types.

## 7 Acknowledgments

This research was undertaken with the assistance of resources from the National Computational Infrastructure (NCI Australia), an NCRIS-enabled capability supported by the Australian Government and the Sydney Informatics Hub, a Core Research Facility of the University of Sydney.

## A Parameter estimates

We model normal epithelial tissue that is just becoming carcinogenic; therefore, we determine our parameter values for cell size, proliferation, migration, and death rate based on data from epithelial cells. Epithelial cells differ in size according to their locations. For instance, the average diameter of a hepatocyte (epithelial cell in the liver) is 18.4 µm, with the majority of hepatocytes’ diameters between 12 and 26 µm [64]. The large squamous epithelial cells and small cuboidal epithelial cells of the lung surface are about 50 µm to 100 µm and 10 µm in diameter, respectively [65]. In addition, the central corneal basal epithelial cell’s diameter is 17.1 *±* 0.8 µm [66]. The human basal epithelial cells of the tongue have a diameter of 10 to 15 µm [67, 68]. From this information, we assume cells in our model have diameters around 15 µm, which is similar to the size of basal epithelial cells of the tongue. We update our model in time steps of length Δ*t* = 1 day.

Researchers estimated that outer limbal basal epithelial cells divide every 8 days, inner limbal basal epithelial cells divide every 4 days, and corneal basal cells divide every 3.5 days [69]. In an *in vitro* study on oral mucosal keratinocytes, Rusu *et al.* showed that the majority of cells in their experiment actively proliferated every 14 days [70], which was similar to results obtained by Klingbeil *et al.*, who showed that the average time for cell proliferation by direct explant technique was 14.2 days [71]. In another *in vitro* analysis on oral mucosal keratinocytes, the cell proliferation time was 9 days [72]. However, Shwetha *et al.* used the direct explant method and found that the average time to cell proliferation ranged from 10 to 12 days [73]. Taking this information into account, we assume each pre-malignant cell (having both *V_G_* and *W_G_* mutations) has a baseline proliferation rate *ρ_V_* _+_*_W_* = 1*/*8 day*^−^*^1^, since *V* + *W* cells have slightly higher replication potential than healthy epithelial cells. In his experiment, Sou found that the *SNAI2* expressing cells’ proliferation rate was almost identical to that of pre-malignant cells, or control cells [23]. Therefore, we assume that the *Y_G_* mutation in the cell does not alter its proliferation profile. So, *Y* cell’s proliferation rate is equal to the pre-malignant cells at *ρ_Y_* = 1*/*8 day*^−^*^1^. Since *X* cells are hyperproliferative and Sou found that these cells’ (*HRAS*-expressing) proliferation rate was twice that of control, or pre-malignant cells [23], we assume the *X_G_* mutation in the cell doubles the proliferation rate from the baseline. Hence, the proliferation rate of *X* cells is *ρ_X_* = *ρ_V_* _+_*_W_* + *d_X_* = 1*/*4 day*^−^*^1^, where *d_X_* = 1*/*8 day*^−^*^1^ is the increase in cell proliferation rate due to the *X_G_* mutation. In addition, he found that the proliferation rate of cells expressing both *HRAS* and *SNAI2* was approximately equal to that of *HRAS*-expressing cells [23]. So, for simplicity, we assume the same proliferation rate *ρ_X_*_+_*_Y_* = *ρ_V_* _+_*_W_* + *d_X_* = 1*/*4 day*^−^*^1^ for *X* + *Y* cells, unless we consider cross-talk interference in the signalling pathway, in which case, we assume *ρ_X_*_+_*_Y_* = *ρ_V_* _+_*_W_* + (1 *− f*)*d_X_* = 1*/*4 day*^−^*^1^, where *f ∈* [0, 1] is an interference factor (see details in Section 2.1).

We assume the minimum distance *r*_min_ required between a cell and its neighbours for it to divide is 0.8 times the average cell size, so *r*_min_ = 0.8 *×* 15 µm = 12 µm. When a cell divides, we assume a new cell centre appears at a distance *d* from the original cell centre, and we assume *d* is 0.05 times the average cell size, so *d* = 0.05 *×* 15 µm = 0.75 µm.

Regarding motility, we assume a pre-malignant *V* +*W* cell has some baseline motility rate at *λ_V_* _+_*_W_* = 0.1 day*^−^*^1^. We choose this value because if *λ_V_* _+_*_W_* is too large or too small, cells move unrealistically fast or slowly, in both cases preventing normal cell division and mutation. Also, *λ_V_* _+_*_W_* = 0.1 day*^−^*^1^ is a reasonable and biologically consistent choice with the other parameters in our model. To keep things simple, we assume the *X_G_*mutation in the cell does not alter motility, maintaining the same motility rate at *λ_X_* = 0.1 day*^−^*^1^ for *X* cells. However, Sou found that *SNAI2*-expressing cells’ motility rate was about double that of control or *HRAS*-expressing cells, and they exhibited nearly the same motility rate as cells expressing both *HRAS* and *SNAI2* [23]. Therefore, we assume the same motility rates for *Y* and *X* +*Y* cells, i.e. *λ_Y_* = *λ_V_* _+_*_W_* +*m_Y_* = 0.2 day*^−^*^1^ and *λ_X_*_+_*_Y_* = *λ_V_* _+_*_W_* +*m_Y_* = 0.2 day*^−^*^1^, where *m_Y_* = 0.1 day*^−^*^1^ is the increase the motility rate due to the *Y_G_*mutation. For the case of cross-talk interference in the signalling pathway, we assume *λ_X_*_+_*_Y_* = *λ_V_* _+_*_W_* + (1 *− f*)*m_Y_* = 0.2 day*^−^*^1^, where *f* is an interference factor as before (see details in Section 2.1). We set *s* to be equal to the average cell size, so *s* = 15 µm. This estimate has reasonable experimental support.

For microenvironmental parameters *r* and *α* in (3) and (4), we assume that *r* is 4-cell diameters, so *r* = 4 *×* 15 µm = 60 µm, and the parameter *α* is 0.25; however, we varied these parameters to investigate their influence.

Meza *et al.* used gestational mutation rates ranging from 10*^−^*^6^ to 10*^−^*^4^ per cell per year for cells at birth [74]. Brash *et al.* used mutation probabilities ranging from 10*^−^*^5^ to 10*^−^*^3^ per gene per cell division for cells affected by UVB irradiation, and 10*^−^*^3^ to 10*^−^*^2^ per gene per cell division for cells affected by UVB radiation [56]. On the other hand, Frank used a mutation probability of 10*^−^*^7^ per gene per cell division for healthy cells and 10*^−^*^3^ per gene per cell division for mutator cells [75]. Based on this information and the fact that *V_a_*+ *W_a_* cells are defective in DNA repair and programmed cell death (akin to a *TP53* mutation), we set their mutation probability, *ε* = 10*^−^*^4^ per gene per cell division.

In addition, cell lifespans also vary across organs. For small intestinal epithelial cells, lifespans range from 3 to 5 days; for colon epithelial cells, they range from 5 to 7 days [76]; and for corneal basal epithelial cells, they are about 7 days [77]. The average turnover of a hepatocyte is from 200 to 400 days [78]. Lifespans of epithelial cells of oral mucosa are usually between 7 and 14 days [79], and lifespans of epithelial cells of the tongue (taste buds) range approximately between 8 and 24 days [80–82]. Premalignant cells and mutant cells have slightly longer lifespans than healthy epithelial cells; therefore, we set our mutant cells’ death rate *r*_death_ = 1*/*16 day*^−^*^1^, which is within that range.

Furthermore, for convenience, we use the same ECM production rate *σ_g_* = 0.01 day*^−^*^1^ as in the VCBM of Lobo [8]. We assume the degradation rate *σ_d_* = 0.0007 cell*^−^*^1^ day*^−^*^1^ to prevent the rapid destruction or disappearance of the underlying ECM. As a starting point, we assume the values for *r_x_* and *r*_deg_ are 3-cell diameters, or 3 *×* 15 µm = 45 µm and 2-cell diameters, or 2 *×* 15 µm = 30 µm, respectively.

For convenience, we begin all models with an initial state consisting of all *V* + *W* cells. In other words, we assume the tissue has developed to a state where it is entirely contained within a pre-malignant field, which is a plausible starting condition before progression to a malignant lesion. These *V* + *W* cells can then acquire *X_G_*or *Y_G_*mutations and give rise to *X* cells or *Y* cells, which can undergo subsequent mutations and produce *X* +*Y* cells. Detailed descriptions and calibrated values of the parameters used in the model are presented in Table A1. We coded and simulated the computational VCBM using MATLAB version R2024b and performed all statistical analyses of the data using GraphPad Prism version 10.4.0. The MATLAB codes to generate the simulations are available on GitHub (https://github.com/Khatun131205/VCBM- of-epithelial-carcinogenesis-evolution).

For clarity, we also present the assumptions and corresponding parameter values used in the four scenarios and the comprehensive model in the form of tables in Appendix B.

**Table A1:**
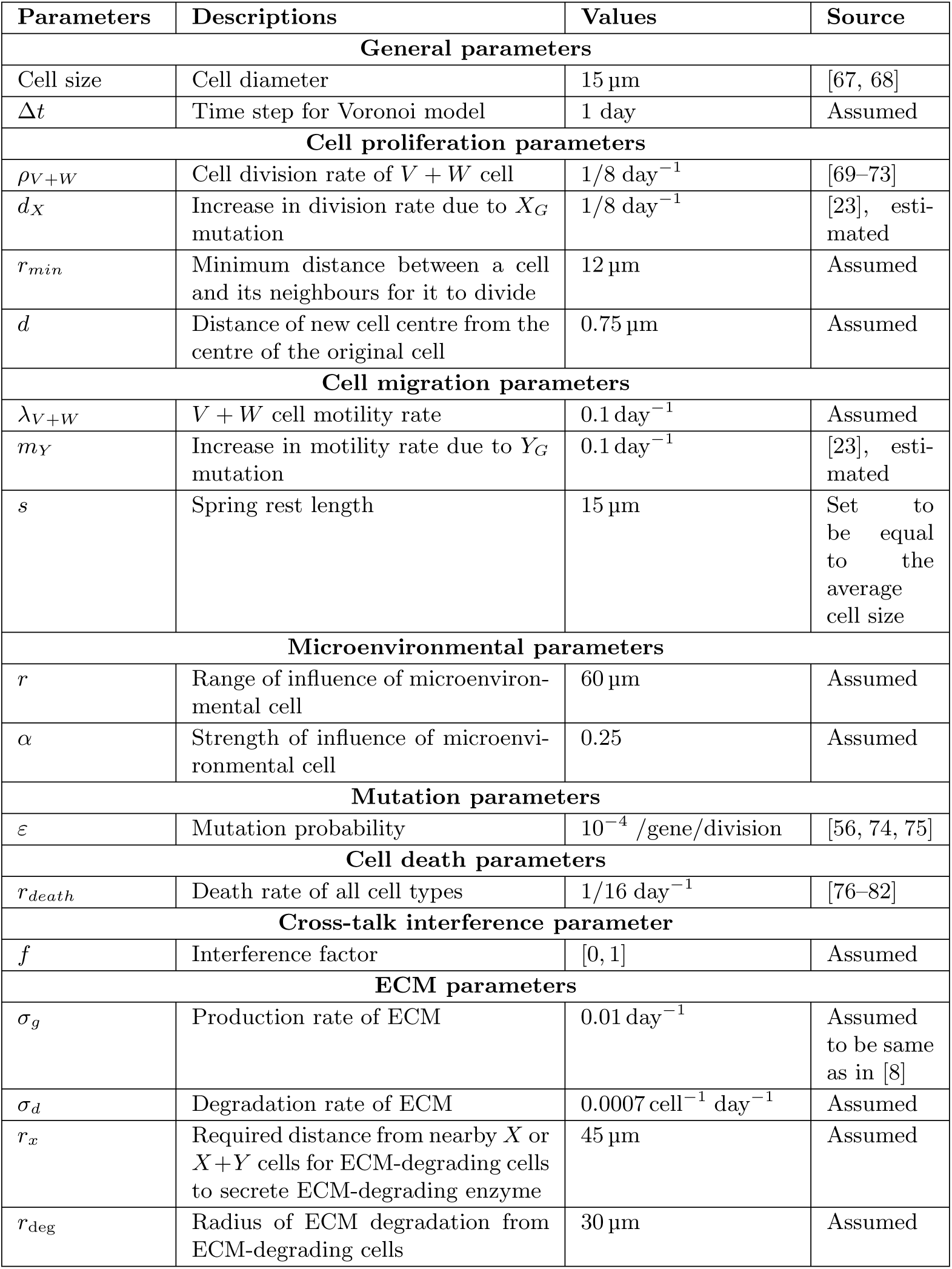
Parameters, their descriptions and default values in the evolutionary model.

| Parameters | Descriptions | Values | Source |
|---|---|---|---|
| **General parameters** |  |  |  |
| Cell size | Cell diameter | 15 µm | [67, 68] |
| Δt | Time step for Voronoi model | 1 day | Assumed |
| **Cell proliferation parameters** |  |  |  |
| ρ<sub>V+W</sub> | Cell division rate of V + W cell | 1/8 day<sup>−1</sup> | [69–73] |
| d<sub>X</sub> | Increase in division rate due to X<sub>G</sub> mutation | 1/8 day<sup>−1</sup> | [23], esti-mated |
| r<sub>min</sub> | Minimum distance between a cell and its neighbours for it to divide | 12 µm | Assumed |
| d | Distance of new cell centre from the centre of the original cell | 0.75 µm | Assumed |
| **Cell migration parameters** |  |  |  |
| λ<sub>V+W</sub> | V + W cell motility rate | 0.1 day<sup>−1</sup> | Assumed |
| m<sub>Y</sub> | Increase in motility rate due to Y<sub>G</sub> mutation | 0.1 day<sup>−1</sup> | [23], esti-mated |
| s | Spring rest length | 15 µm | Set to be equal to the average cell size |
| **Microenvironmental parameters** |  |  |  |
| r | Range of influence of microenviron-mental cell | 60 µm | Assumed |
| α | Strength of influence of microenvi-ronmental cell | 0.25 | Assumed |
| **Mutation parameters** |  |  |  |
| ε | Mutation probability | 10<sup>−4</sup> /gene/division | [56, 74, 75] |
| **Cell death parameters** |  |  |  |
| r<sub>death</sub> | Death rate of all cell types | 1/16 day<sup>−1</sup> | [76–82] |
| **Cross-talk interference parameter** |  |  |  |
| f | Interference factor | [0, 1] | Assumed |
| **ECM parameters** |  |  |  |
| σ<sub>g</sub> | Production rate of ECM | 0.01 day<sup>−1</sup> | Assumed to be same as in [8] |
| σ<sub>d</sub> | Degradation rate of ECM | 0.0007 cell<sup>−1</sup> day<sup>−1</sup> | Assumed |
| r<sub>x</sub> | Required distance from nearby X or X+Y cells for ECM-degrading cells to secrete ECM-degrading enzyme | 45 µm | Assumed |
| r<sub>deg</sub> | Radius of ECM degradation from ECM-degrading cells | 30 µm | Assumed |

## B Proliferation, motility and degradation parameters for each simulation

In this section, we present the model assumptions (discussed in Section 2) and corresponding parameter values used in the simulations of the four scenarios (Scenarios XaYa, XaYm, XmYa and XmYm) in Section 3.1 and the comprehensive model in Section 3.2 in tabular form.

For *ϕ ∈ {a, m}*, the intrinsic proliferation rate is 1*/*8 day*^−^*^1^ for *V_a_* + *W_a_*and *Y_ϕ_*and 1*/*4 day*^−^*^1^ (maximal proliferation rate) for *X_ϕ_*and *X_ϕ_* + *Y_ϕ_*cells. The proliferation rate of *V_a_* + *W_a_* and *Y_ϕ_* cells remains the same when they interact with *X_a_* or *X_a_* + *Y_ϕ_*cells, or with themselves (see Tables B1 and B2); however, their proliferation rates increase when they interact with *X_m_*or *X_m_* + *Y_ϕ_*cells (see Tables B3 and B4).

In addition, for *V_a_* + *W_a_* and *X_ϕ_* cells, the intrinsic motility rate is 0.1 day*^−^*^1^ and for *Y_ϕ_* and *X_ϕ_* + *Y_ϕ_* cells, it is 0.2 day*^−^*^1^ (maximal motility rate). The motility rate of *V_a_* + *W_a_* and *X_ϕ_* cells remains the same when they interact with *Y_a_* or *X_ϕ_*+ *Y_a_* cells, or with themselves (see Tables B1 and B3), but their motility rates increase when they interact with *Y_m_* or *X_ϕ_*+ *Y_m_* cells (see Tables B2 and B4).

Furthermore, for *V_a_* + *W_a_*, *X_ϕ_*, and *Y_ϕ_* cells, the intrinsic degradation rate is 0 cell*^−^*^1^day*^−^*^1^ and for *X_a_* + *Y_ϕ_* cells, it is 0.0007 cell*^−^*^1^day*^−^*^1^. *V_a_* + *W_a_* and *X_ϕ_*cells do not secrete ECM-degrading enzymes, so the degradation rate of these cells remains 0. However, *Y_ϕ_* cells’ degradation rate becomes 0.0007 cell*^−^*^1^day*^−^*^1^ when they interact with *X_a_* or *X_a_*+ *Y_ϕ_* cells (see Tables B1 and B2). The degradation rate of *Y_ϕ_* cells increases to 0.0009 cell*^−^*^1^day*^−^*^1^ when they interact with *X_m_*or *X_m_* + *Y_ϕ_*cells and *X_m_* + *Y_ϕ_*cells’ degradation rate also rises to 0.0009 cell*^−^*^1^day*^−^*^1^ as they have *X_m_* mutation with themselves (see Tables B3 and B4).

**Table B1:**
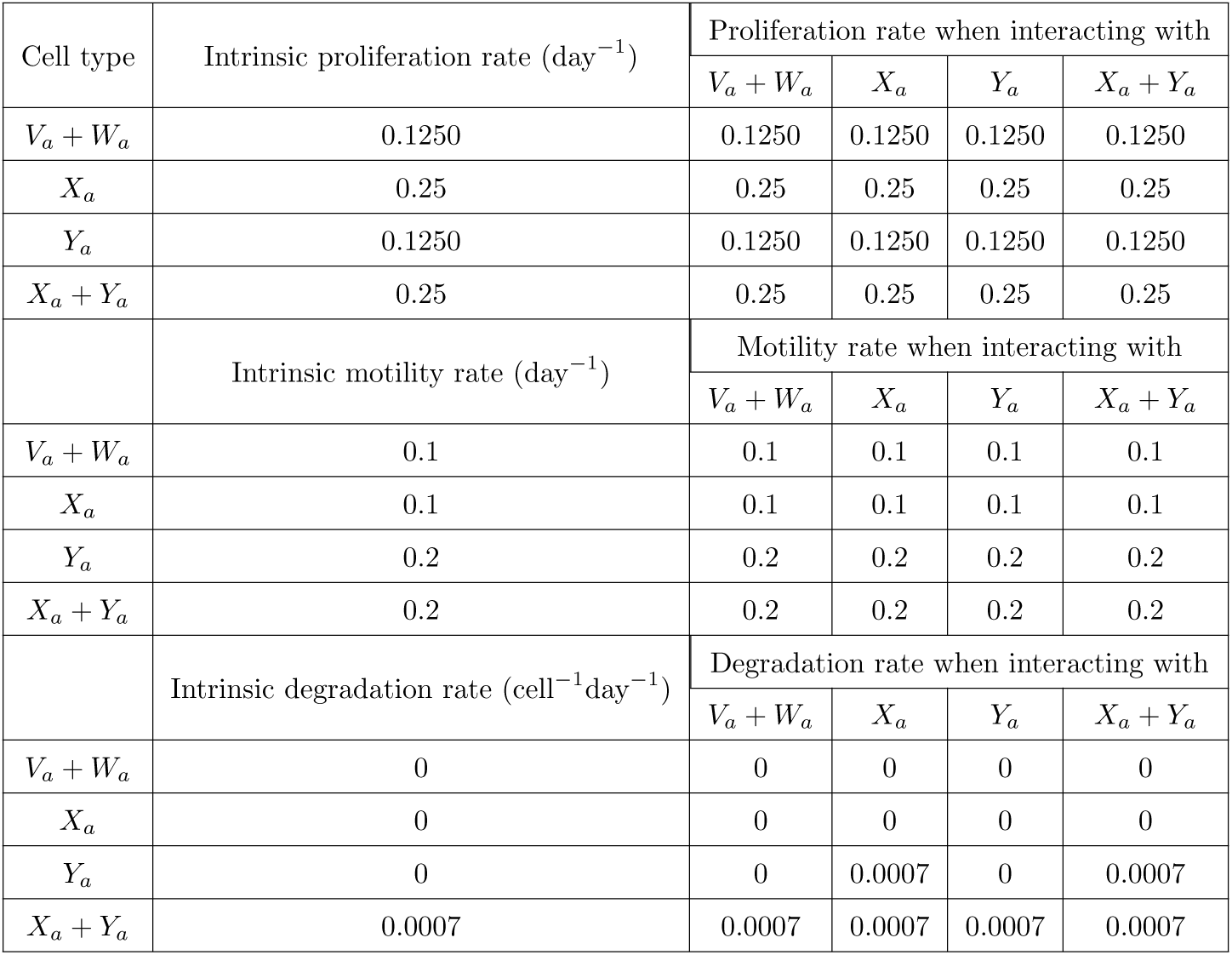
Cell proliferation, motility and ECM degradation rates in Scenario XaYa.

**Table B2:**
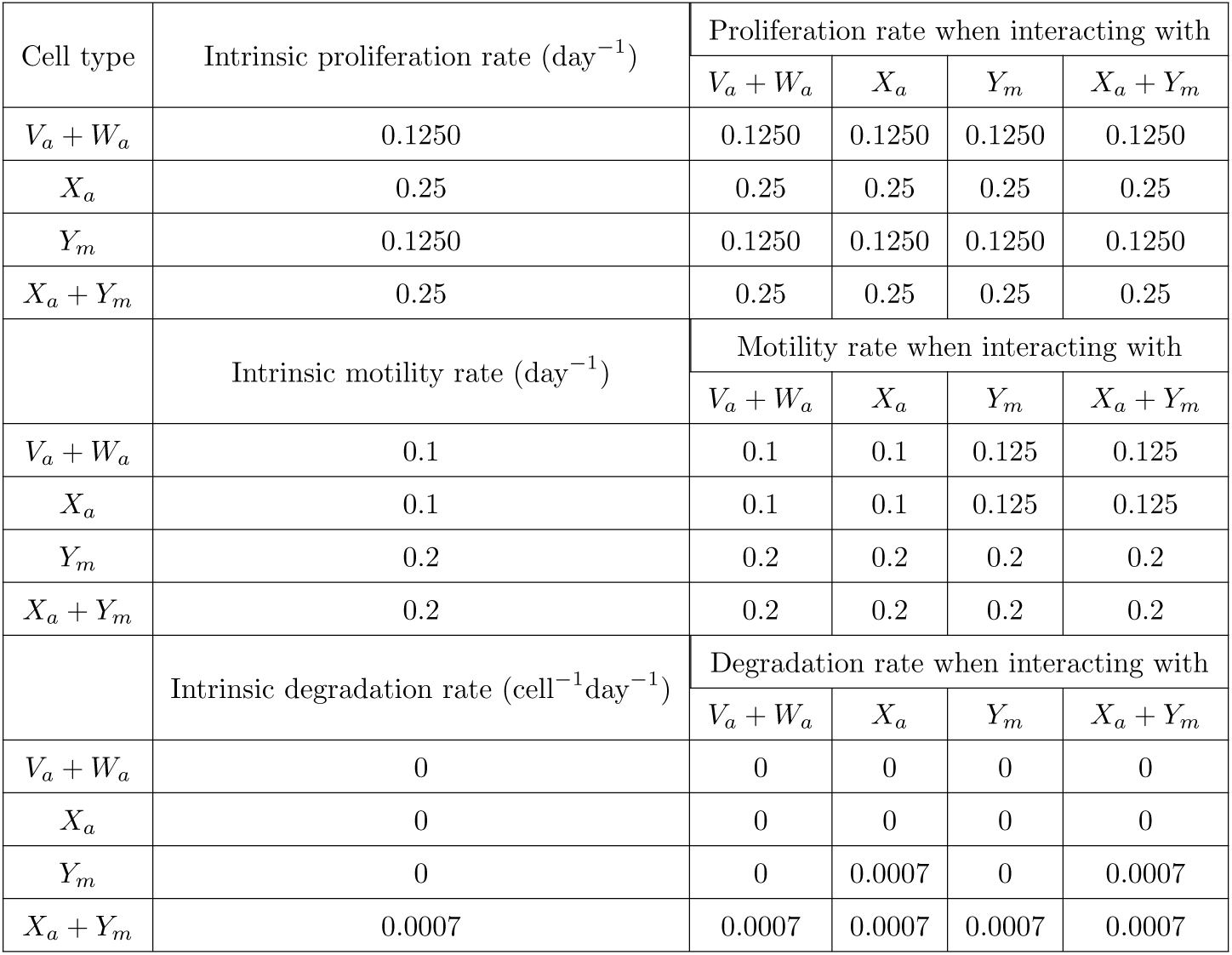
Cell proliferation, motility and ECM degradation rates in Scenario XaYm.

| Cell type | Intrinsic proliferation rate (day<sup>−1</sup>) | Proliferation rate when interacting with |  |  |  |
|  |  | *V<sub>a</sub>* + *W<sub>a</sub>* | *X<sub>a</sub>* | *Y<sub>m</sub>* | *X<sub>a</sub>* + *Y<sub>m</sub>* |
|---|---|---|---|---|---|
| *V<sub>a</sub>* + *W<sub>a</sub>* | 0.1250 | 0.1250 | 0.1250 | 0.1250 | 0.1250 |
| *X<sub>a</sub>* | 0.25 | 0.25 | 0.25 | 0.25 | 0.25 |
| *Y<sub>m</sub>* | 0.1250 | 0.1250 | 0.1250 | 0.1250 | 0.1250 |
| *X<sub>a</sub>* + *Y<sub>m</sub>* | 0.25 | 0.25 | 0.25 | 0.25 | 0.25 |
|  | Intrinsic motility rate (day<sup>−1</sup>) | Motility rate when interacting with |  |  |  |
|  |  | *V<sub>a</sub>* + *W<sub>a</sub>* | *X<sub>a</sub>* | *Y<sub>m</sub>* | *X<sub>a</sub>* + *Y<sub>m</sub>* |
| *V<sub>a</sub>* + *W<sub>a</sub>* | 0.1 | 0.1 | 0.1 | 0.125 | 0.125 |
| *X<sub>a</sub>* | 0.1 | 0.1 | 0.1 | 0.125 | 0.125 |
| *Y<sub>m</sub>* | 0.2 | 0.2 | 0.2 | 0.2 | 0.2 |
| *X<sub>a</sub>* + *Y<sub>m</sub>* | 0.2 | 0.2 | 0.2 | 0.2 | 0.2 |
|  | Intrinsic degradation rate (cell<sup>−1</sup>day<sup>−1</sup>) | Degradation rate when interacting with |  |  |  |
|  |  | *V<sub>a</sub>* + *W<sub>a</sub>* | *X<sub>a</sub>* | *Y<sub>m</sub>* | *X<sub>a</sub>* + *Y<sub>m</sub>* |
| *V<sub>a</sub>* + *W<sub>a</sub>* | 0 | 0 | 0 | 0 | 0 |
| *X<sub>a</sub>* | 0 | 0 | 0 | 0 | 0 |
| *Y<sub>m</sub>* | 0 | 0 | 0.0007 | 0 | 0.0007 |
| *X<sub>a</sub>* + *Y<sub>m</sub>* | 0.0007 | 0.0007 | 0.0007 | 0.0007 | 0.0007 |

**Table B3:** Cell proliferation, motility and ECM degradation rates in Scenario XmYa.

| Cell type | Intrinsic proliferation rate (day<sup>−1</sup>) | Proliferation rate when interacting with |  |  |  |
|---|---|---|---|---|---|
|  |  | *V<sub>a</sub>* + *W<sub>a</sub>* | *X<sub>m</sub>* | *Y<sub>a</sub>* | *X<sub>m</sub>* + *Y<sub>a</sub>* |
| *V<sub>a</sub>* + *W<sub>a</sub>* | 0.1250 | 0.1250 | 0.1562 | 0.1250 | 0.1562 |
| *X<sub>m</sub>* | 0.25 | 0.25 | 0.25 | 0.25 | 0.25 |
| *Y<sub>a</sub>* | 0.1250 | 0.1250 | 0.1562 | 0.1250 | 0.1562 |
| *X<sub>m</sub>* + *Y<sub>a</sub>* | 0.25 | 0.25 | 0.25 | 0.25 | 0.25 |
|  | Intrinsic motility rate (day<sup>−1</sup>) | Motility rate when interacting with |  |  |  |
|  |  | *V<sub>a</sub>* + *W<sub>a</sub>* | *X<sub>m</sub>* | *Y<sub>a</sub>* | *X<sub>m</sub>* + *Y<sub>a</sub>* |
| *V<sub>a</sub>* + *W<sub>a</sub>* | 0.1 | 0.1 | 0.1 | 0.1 | 0.1 |
| *X<sub>m</sub>* | 0.1 | 0.1 | 0.1 | 0.1 | 0.1 |
| *Y<sub>a</sub>* | 0.2 | 0.2 | 0.2 | 0.2 | 0.2 |
| *X<sub>m</sub>* + *Y<sub>a</sub>* | 0.2 | 0.2 | 0.2 | 0.2 | 0.2 |
|  | Intrinsic degradation rate (cell<sup>−1</sup>day<sup>−1</sup>) | Degradation rate when interacting with |  |  |  |
|  |  | *V<sub>a</sub>* + *W<sub>a</sub>* | *X<sub>m</sub>* | *Y<sub>a</sub>* | *X<sub>m</sub>* + *Y<sub>a</sub>* |
| *V<sub>a</sub>* + *W<sub>a</sub>* | 0 | 0 | 0 | 0 | 0 |
| *X<sub>m</sub>* | 0 | 0 | 0 | 0 | 0 |
| *Y<sub>a</sub>* | 0 | 0 | 0.0009 | 0 | 0.0009 |
| *X<sub>m</sub>* + *Y<sub>a</sub>* | 0.0009 | 0.0009 | 0.0009 | 0.0009 | 0.0009 |

**Table B4:**
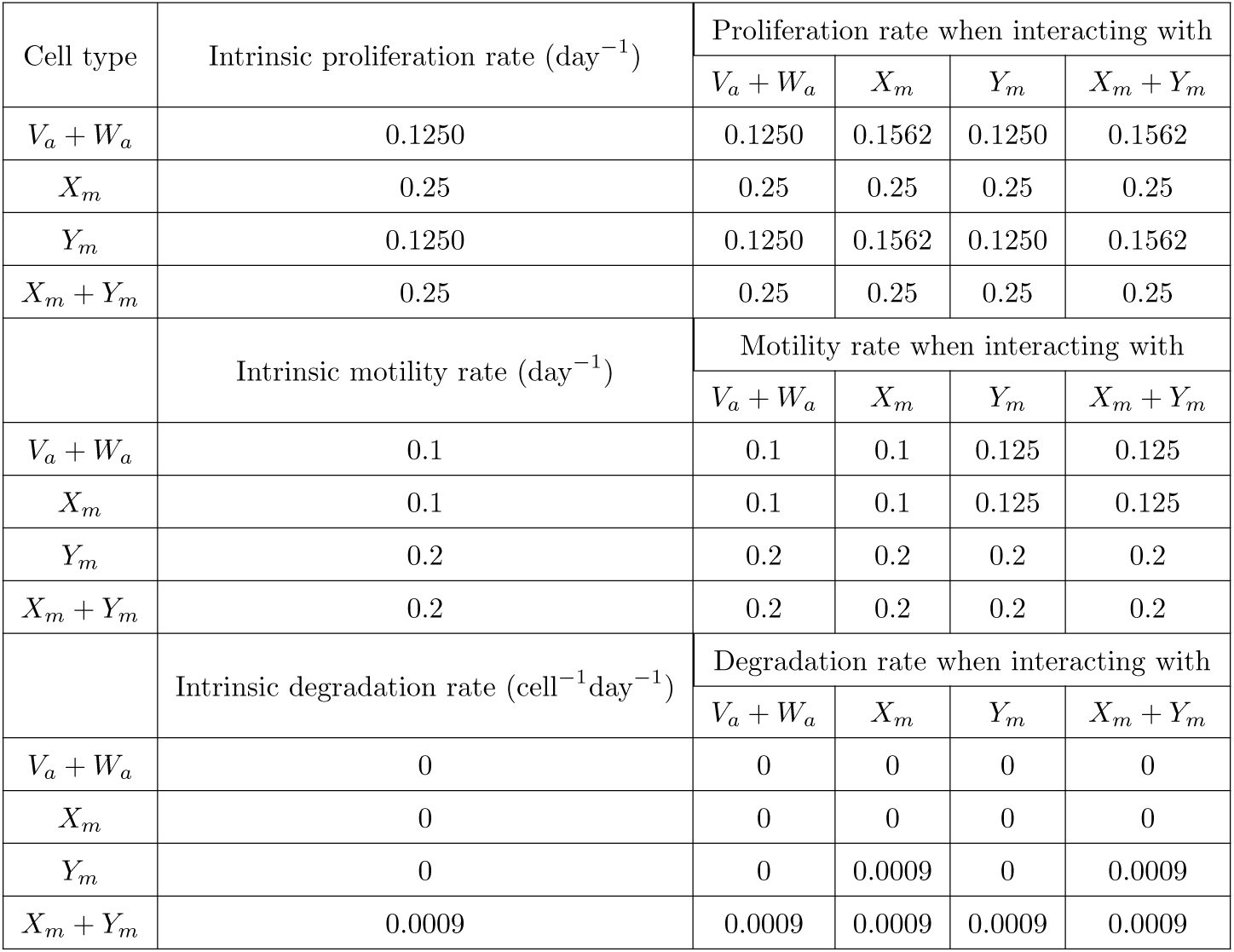
Cell proliferation, motility and ECM degradation rates in Scenario XmYm.

**Table B5:**
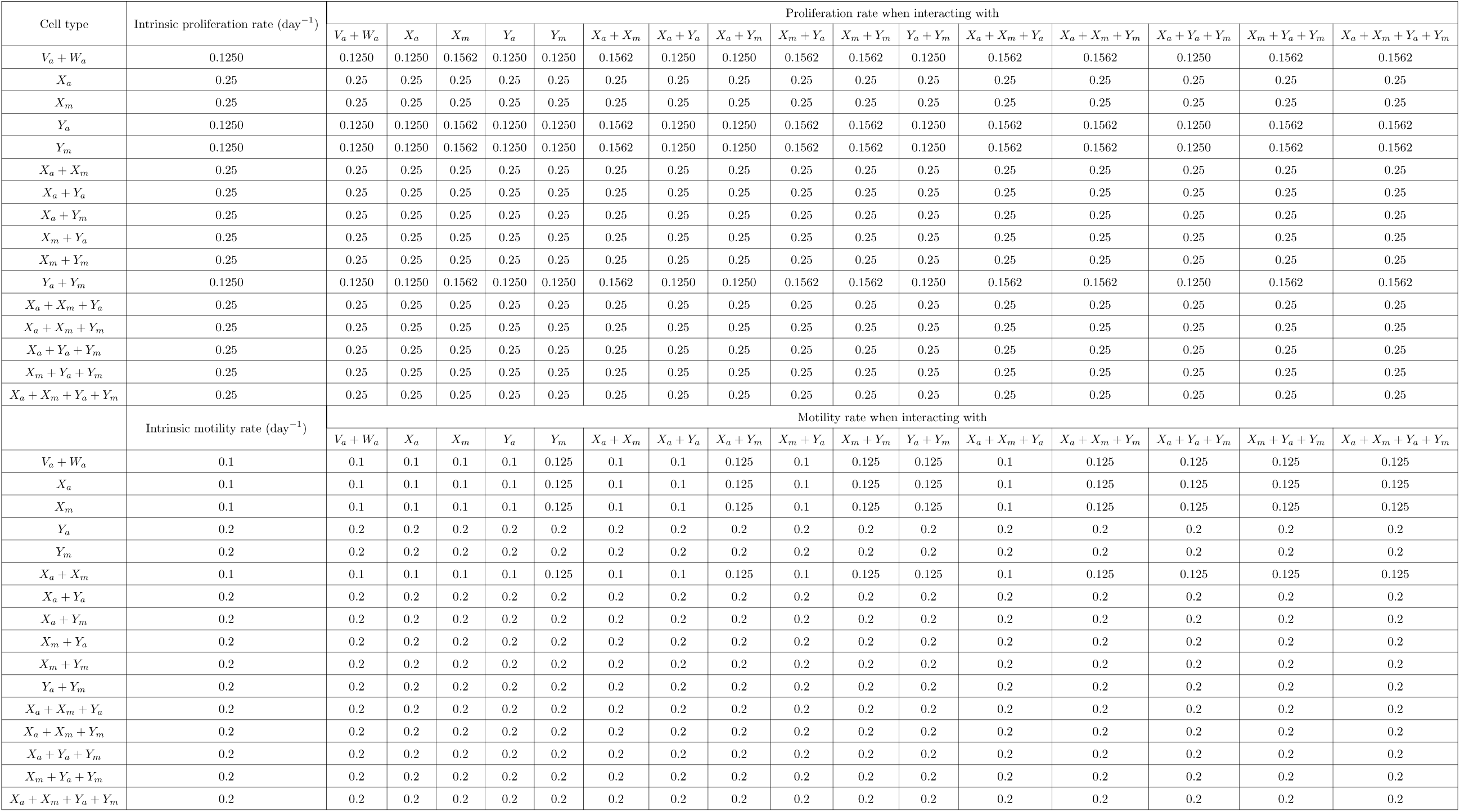
Cell proliferation and motility rates in the comprehensive model.

## C More Visual representation of outcomes

In this section, we illustrate the evolution of carcinogenic lesions via the Scenarios XaYa, XaYm, XmYa and XmYm, as well as the comprehensive model.

**Figure C1:**
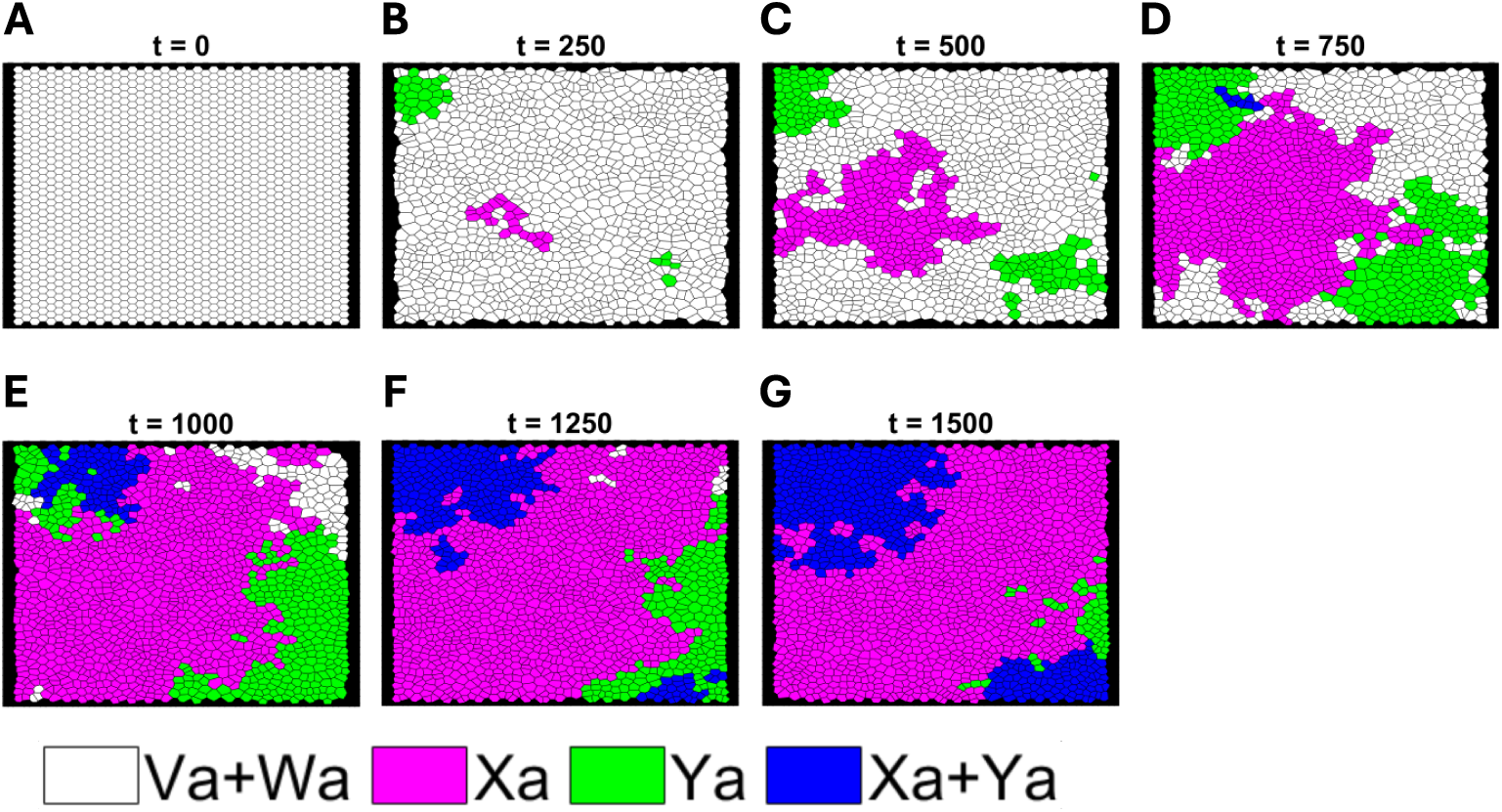
A representative carcinogenic lesion progression in Scenario XaYa. The seven snapshots represent the Voronoi tessellations at times *t* = 0, 250, 500, 750, 1000, 1250 and 1500 days (**A-G**).

**Figure C2:**
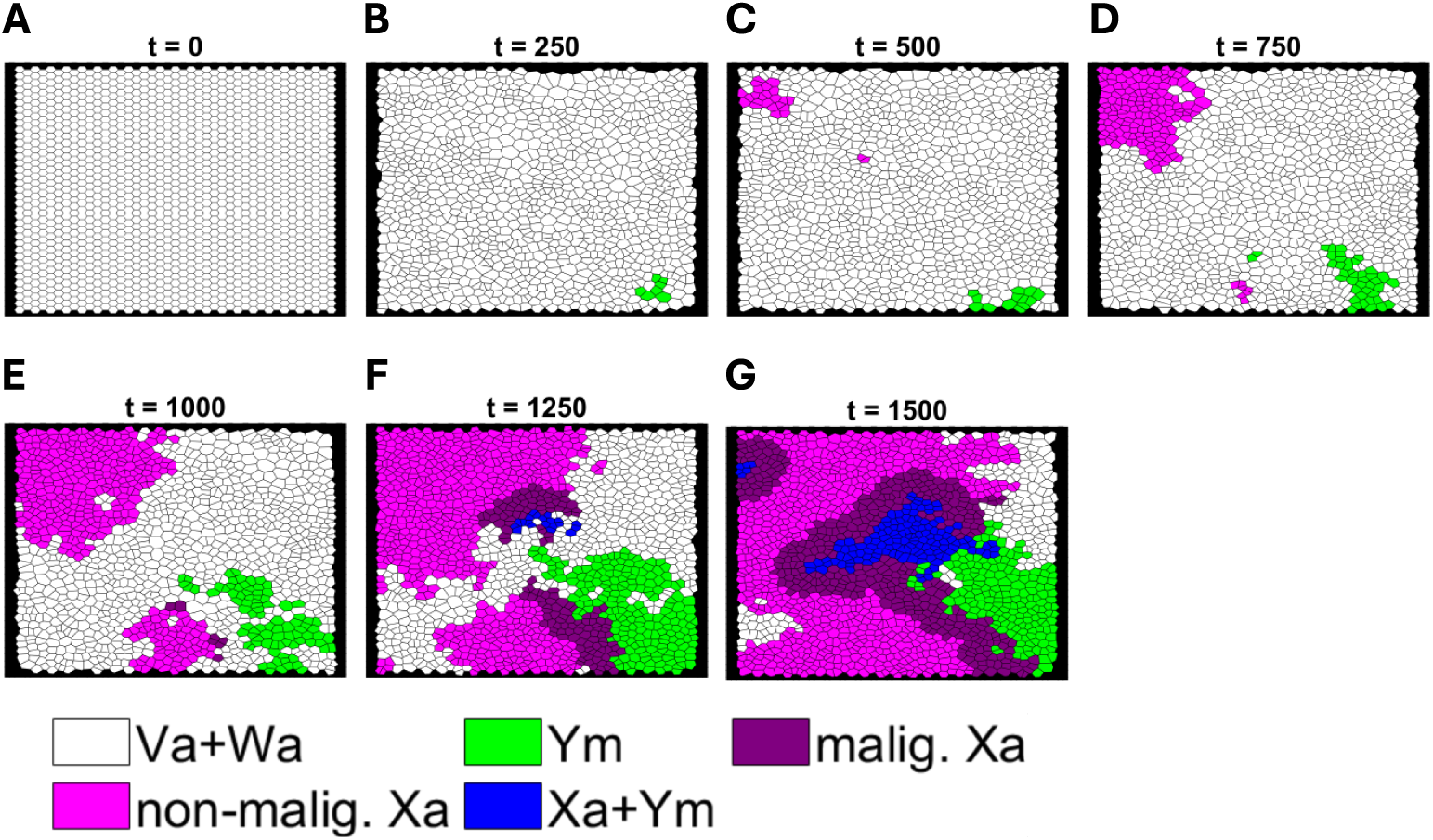
A representative carcinogenic lesion progression in Scenario XaYm. The seven snapshots represent the Voronoi tessellations at times *t* = 0, 250, 500, 750, 1000, 1250 and 1500 days (**A-G**).

**Figure C3:**
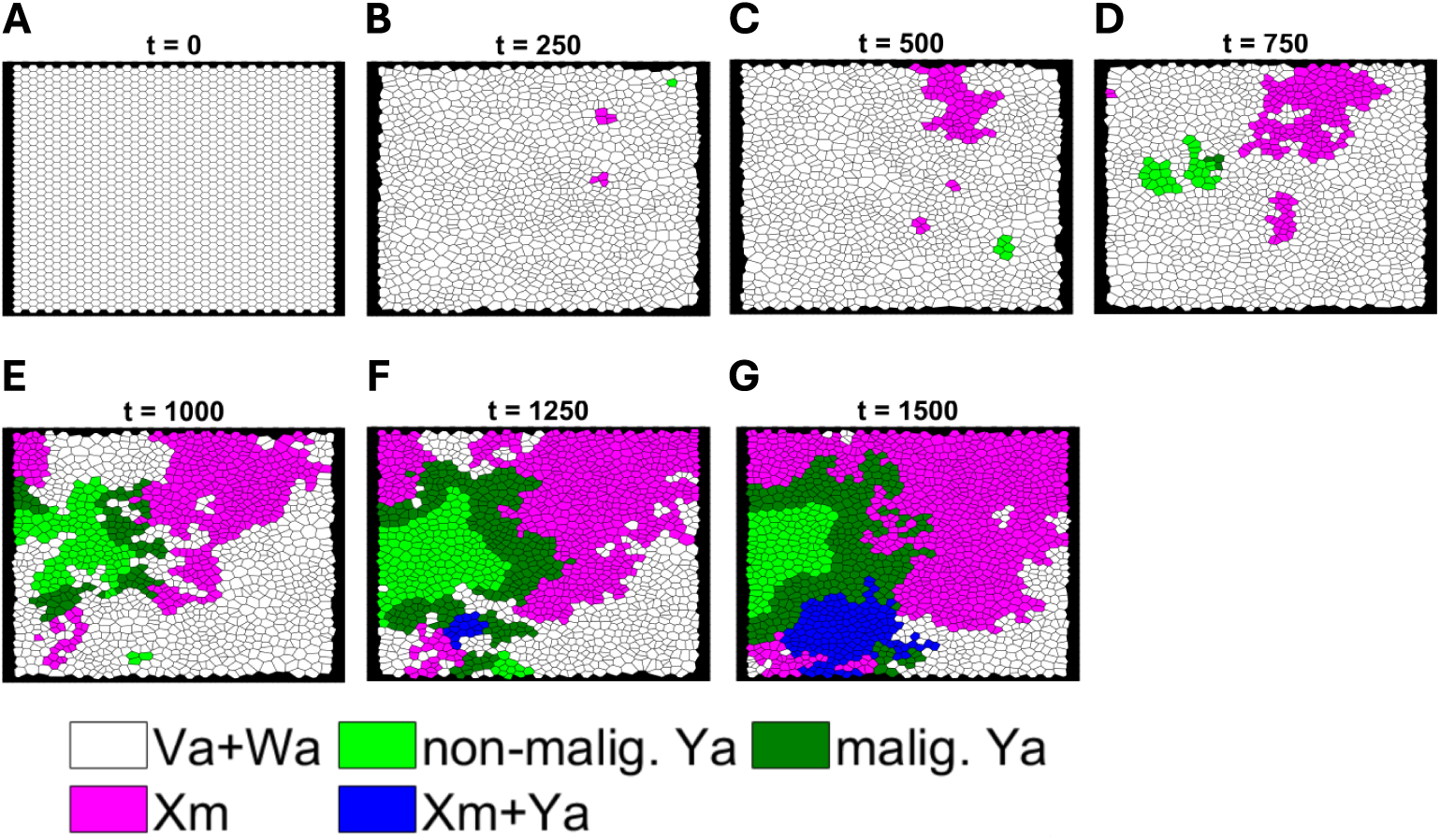
A representative carcinogenic lesion progression in Scenario XmYa. The seven snapshots represent the Voronoi tessellations at times *t* = 0, 250, 500, 750, 1000, 1250 and 1500 days (**A-G**).

**Figure C4:**
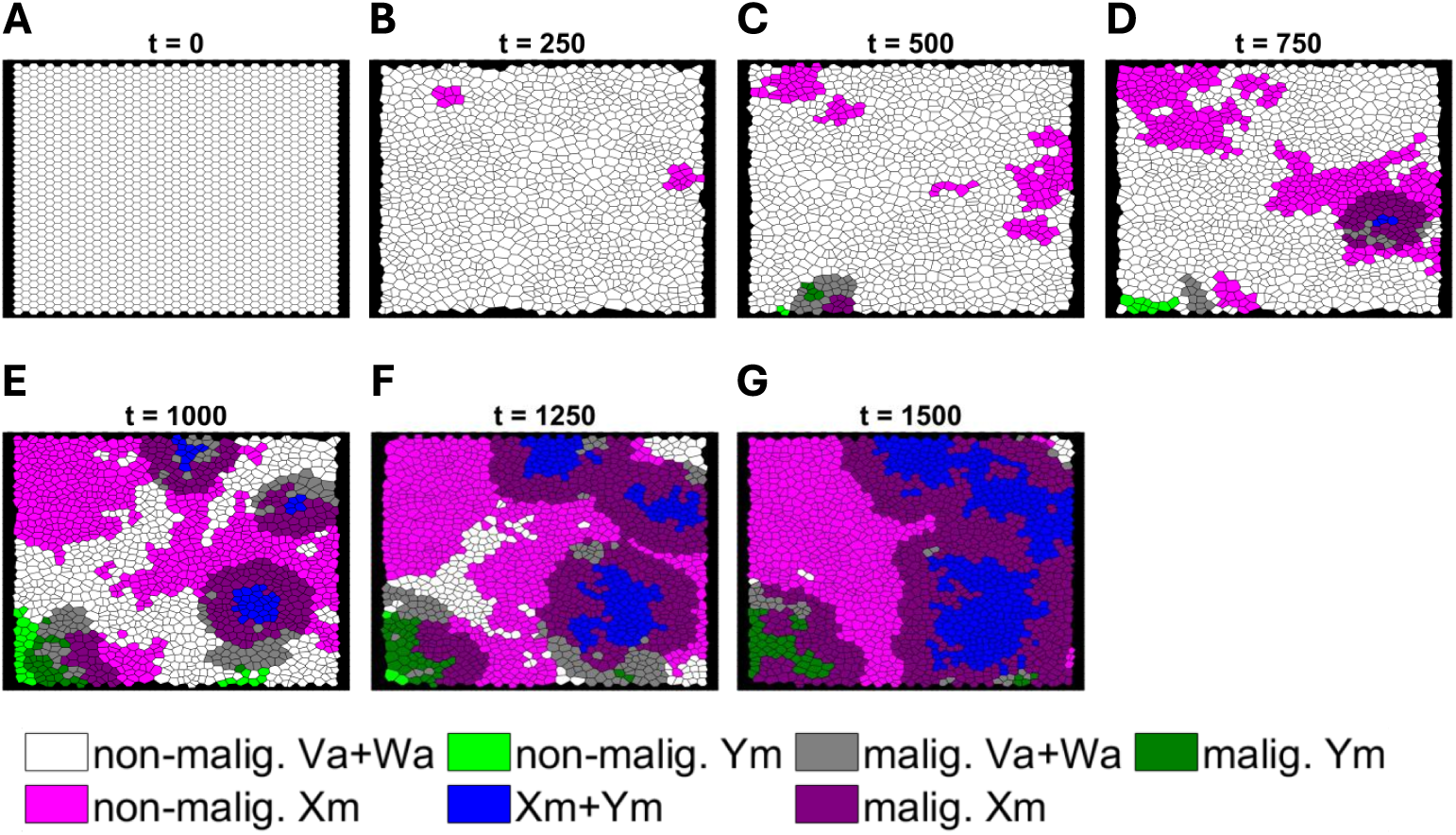
A representative carcinogenic lesion progression in Scenario XmYm. The seven snapshots represent the Voronoi tessellations at times *t* = 0, 250, 500, 750, 1000, 1250 and 1500 days (**A-G**).

**Figure C5:**
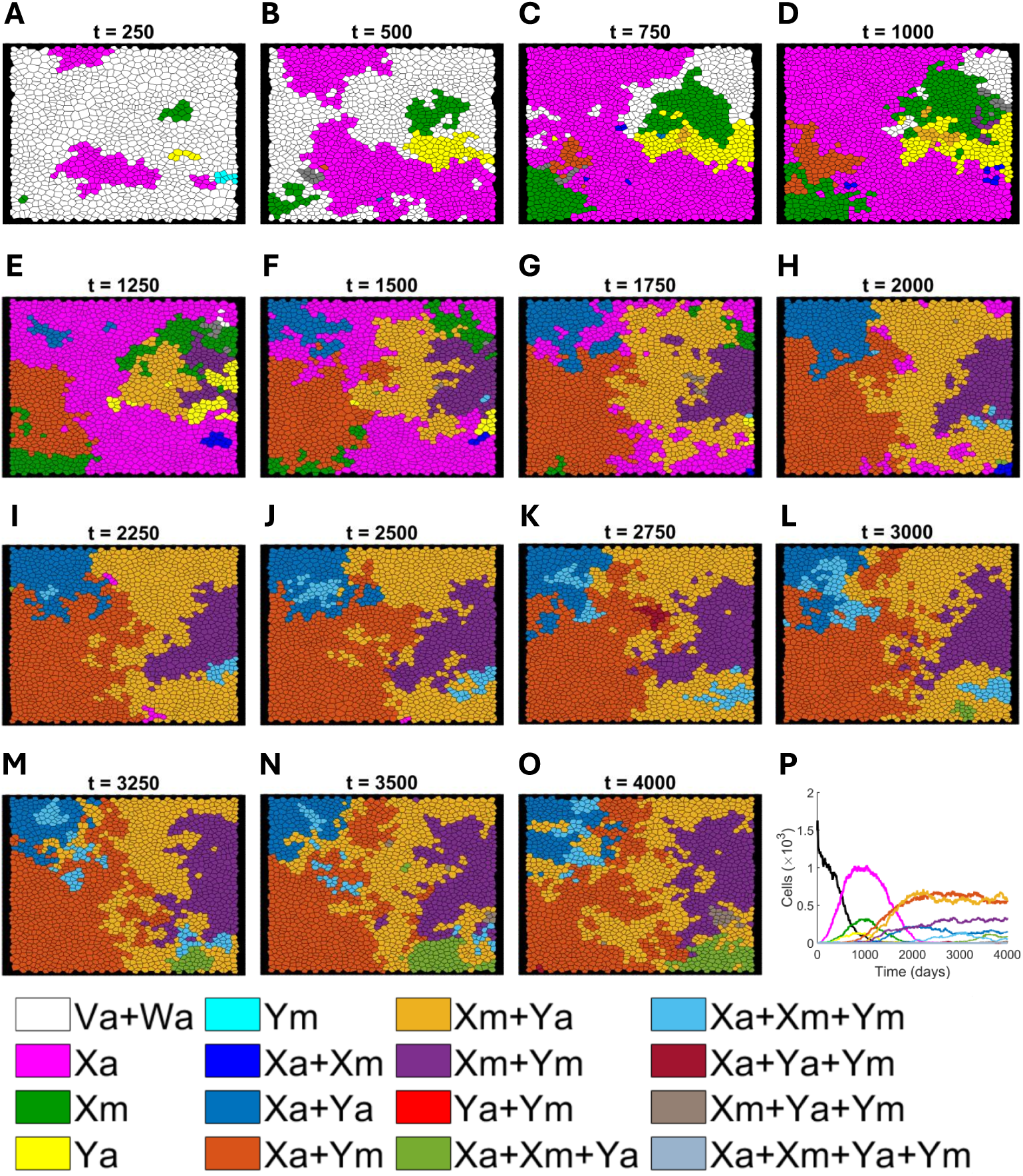
A representative carcinogenic lesion progression in the comprehensive model. The sixteen snapshots represent the Voronoi diagrams at different times *t* (**A-O**), and the corresponding time evolution (**P**).

## D Additional results

In this Section, we present the figures associated with the results discussed in Section 3. These figures are included to support the main findings and to avoid repetition of the detailed explanation in the main text.

**Figure D1:**
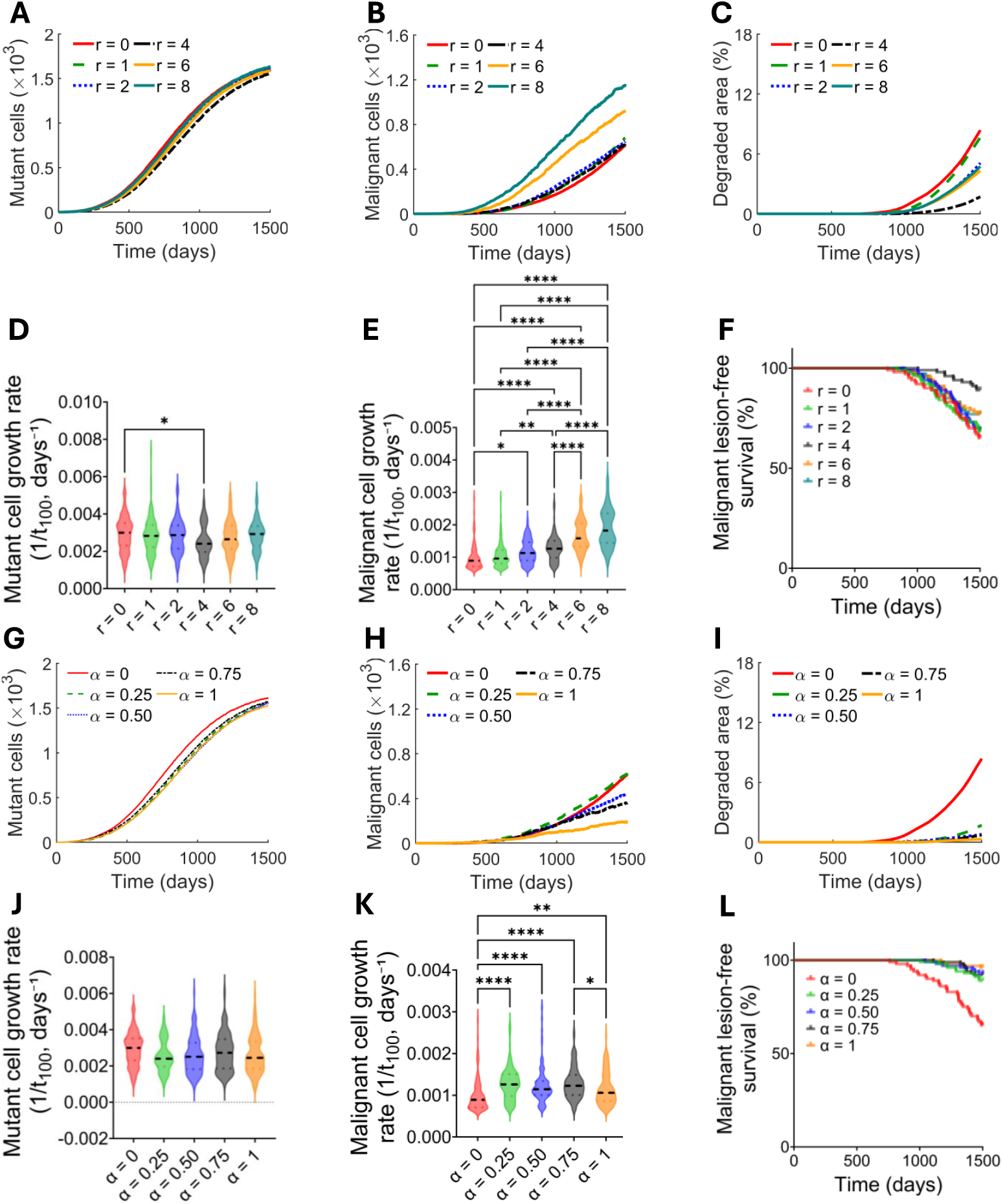
Clonal evolution in Scenario XaYm of epithelial carcinogenesis. The effects of varying the ranges **(A-F)** and strengths **(G-L)** of microenvironmental phenotypes on malignant characteristics are shown for Scenario XaYm. The mean number of mutant cells (**A** and **G**), number of malignant cells (**B** and **H**), and areas of degraded ECM (**C** and **I**) from over 100 simulations over time are shown. The speeds of acquiring malignant characteristics are represented by the inverse of the time elapsed to reach 100 mutant cells (**D** and **J**), 100 malignant cells (**E** and **K**) or by the time of first malignant lesion formation (**F** and **L**). The p-values for differences were determined using the Kruskal-Wallis test followed by Dunn’s multiple comparisons correction. Only pairwise comparisons with *p ≤* 0.05 are shown by asterisks.

**Figure D2:**
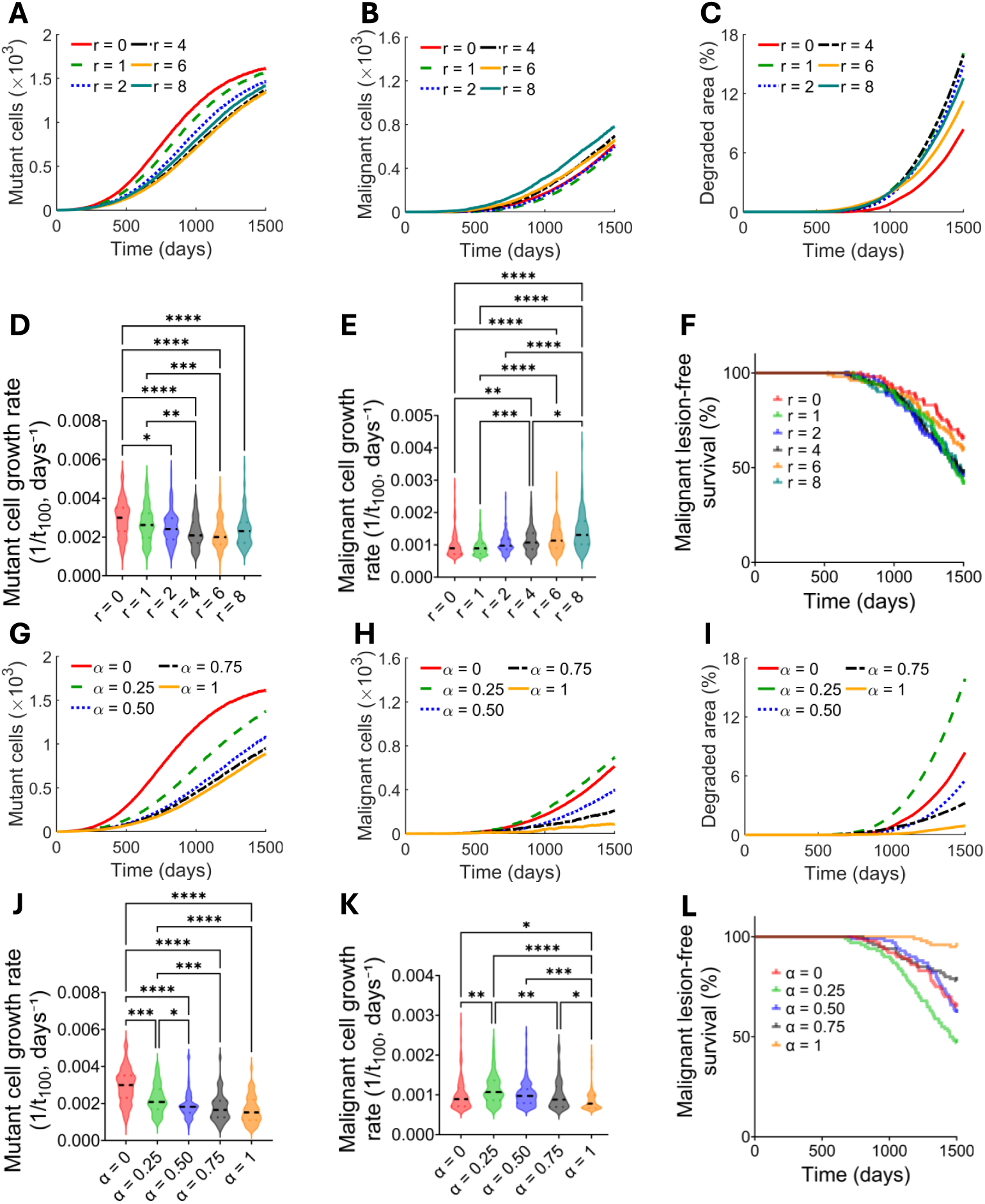
Clonal evolution in Scenario XmYa of epithelial carcinogenesis. The effects of varying the ranges **(A-F)** and strengths **(G-L)** of microenvironmental phenotypes on malignant characteristics are shown for Scenario XmYa. The mean number of mutant cells (**A** and **G**), number of malignant cells (**B** and **H**), and areas of degraded ECM (**C** and **I**) from over 100 simulations over time are shown. The speeds of acquiring malignant characteristics are represented by the inverse of the time elapsed to reach 100 mutant cells (**D** and **J**), 100 malignant cells (**E** and **K**) or by the time of first malignant lesion formation (**F** and **L**). The p-values for differences were determined using the Kruskal-Wallis test followed by Dunn’s multiple comparisons correction. Only pairwise comparisons with *p ≤* 0.05 are shown by asterisks.

**Figure D3:**
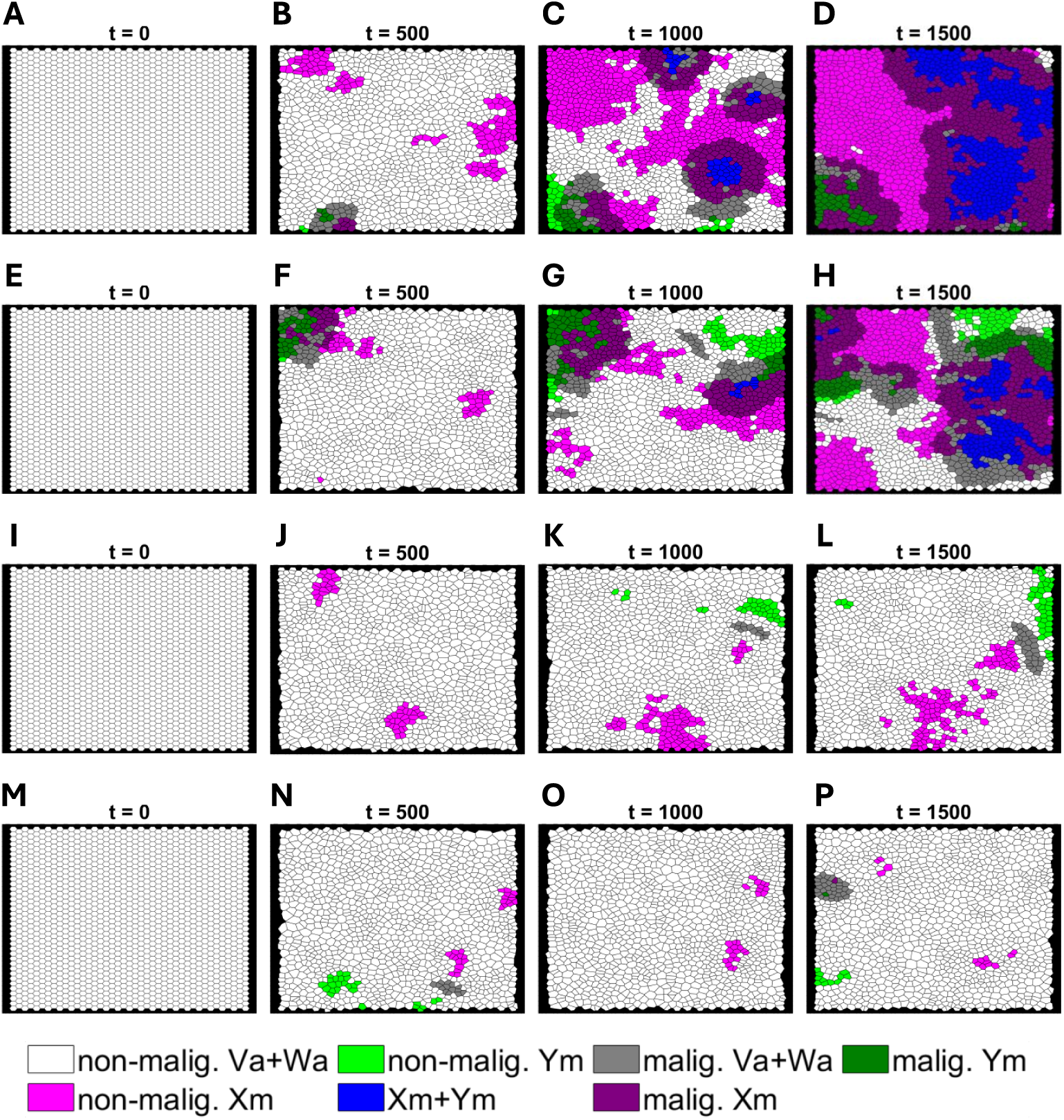
Representative carcinogenic lesion evolution via Scenario XmYm. The sixteen snapshots show carcinogenic lesion growth at *t* = 0, 500, 1000 and 1500 days for four different microenvironmental strengths: **(A-D)** *α* = 0.25, **(E-H)** *α* = 0.50, **(I-L)** *α* = 0.75, and **(M-P)** *α* = 1. These snapshots provide visual evidence of the structural differences for the different microenvironmental strengths from the simulations in Figure 8**(H-M)**.

